# Cryo-EM analysis of the *Bacillus thuringiensis* extrasporal matrix identifies F-ENA as a widespread family of endospore appendages across the Firmicutes phylum

**DOI:** 10.1101/2025.02.11.637640

**Authors:** Mike Sleutel, Adrià Sogues, Nani Van Gerven, Unni Lise Jonsmoen, Inge Van Molle, Marcus Fislage, Laurent Dirk Theunissen, Nathan F. Bellis, Diana P. Baquero, Edward H. Egelman, Mart Krupovic, Fengbin Wang, Marina Aspholm, Han Remaut

## Abstract

For over 100 years, *Bacillus thuringiensis* (Bt) has been used as an agricultural biopesticide to control pests caused by insect species in the orders of Lepidoptera, Diptera and Coleoptera. Under nutrient starvation, Bt cells differentiate into spores and associated toxin crystals that can adopt biofilm-like aggregates. We reveal that such Bt spore/toxin biofilms are embedded in a fibrous extrasporal matrix (ESM), and using cryoID, we resolved the structure and molecular identity of an uncharacterized type of pili, referred to here as <u>F</u>ibrillar <u>EN</u>dospore <u>A</u>ppendages or ‘F-ENA’. F-ENA are monomolecular protein polymers tethered to the exosporium of Bt and are decorated with a flexible tip fibrillum. Phylogenetic analysis reveals that F-ENA is widespread not only in the class Bacilli, but also in the class Clostridia, and the cryoEM structures of F-ENA filaments from *Bacillus, Anaerovorax* and *Paenibaccilus* reveal subunits with a generic head-neck domain structure, where the β-barrel neck of variable length latch onto a preceding head domain through short N-terminal hook peptides. In *Bacillus*, two collagen-like proteins (CLP) respectively tether F-ENA to the exosporium (F-Anchor), or constitute the tip fibrillum at the distal terminus of F-ENA (F-BclA). Sedimentation assays point towards F-ENA involvement in spore-spore clustering, likely mediated via F-BclA contacts and F-ENA bundling through the antiparallel interlocking of the head-neck units.

## Introduction

*Bacillus thuringiensis* (Bt) is a Gram-positive, spore-forming bacterium that occurs among others in soils, on leaf surfaces and in the gut of caterpillars. During sporulation, many Bt strains produce crystalline protein condensates or parasporal bodies (PSBs) with insecticidal activity, known as Cry toxins. These toxins target a diverse range of insect species primarily of the order Lepidoptera (butterflies and moths), Diptera (flies and mosquitoes), Coleoptera (beetles and weevils), and hymenopterans (wasps and bees)^1,2^. When susceptible larvae ingest the Bt toxin crystals, digestive enzymes present in their highly alkaline gut activate the Cry proteins. Binding to specific receptors on the membranes of mid-gut epithelial cells results in pore formation, followed by paralysis and/or rupture of the digestive tract^3,4^. In contrast to poisonous insecticides that target the nervous system, Bt acts by starving infected larvae as they will stop feeding within hours after ingestion. Organisms lacking the appropriate receptors in their gut are not affected by the Cry protein, making each Cry toxin only effective against a specific group of insects^5^.

Spores and parasporal bodies (PSBs) produced by Bt have been used in agriculture to control insect pests since the 1920s^6^. Today, numerous Bt strains producing different toxins are commercially available (e.g. trade names such as Monterey Bt, Milky spore, Mosquito dunks, DiPel and Thuricide). These Bt-based products are regarded as environment-friendly biopesticides because of their high species specificity, with little or no effect on humans, wildlife and pollinators^2,7,8^. However, whether applied as a spray or, less often, as granules, cry toxins biodegrade quickly in sunlight, hence most formulations persist on foliage less than a week following application, and reapplication is often necessary under heavy insect pressure.

Bt has the potential to form biofilms - complex microbial ecosystems embedded in a self-produced matrix of extracellular polymeric substance - that are considered to be a key factor for survival when bacteria face environmental chemical, physical and mechanical stresses^9–11^. Within the biofilm, Bt spores are engulfed by a proteinaceous parasporal sacculus known as the exosporium^12^. The surface of the exosporium is adorned by a dense array of BclAs (bacterial collagen like protein of Anthracis), collectively referred to as the hairy nap^13^. From this nap, fibrous structures termed endospore appendages (ENAs) emanate, contributing to the unique architecture and functionality of the biofilm^14,15^.

In this contribution we sought to further understand the molecular architecture of a Btk spore biofilm. Using cryo-electron transmission electron microscopy (cryoEM) we show that Btk spores are decorated with S-ENA fibers^14^ and a novel type of pili of 5 nm diameter, referred to as F(fibrillar)-type endospore appendages (F-ENA), that are tethered to the exosporium and decorated with a 3 nm diameter tip fibrillum (ruffle) at their spore-distal terminus. We solve the helical fiber ultrastructure and show that the major subunit is a 10 kDa protein that belongs to the DUF4183 family. Using an integrative structural biology approach combining cryoEM, AlphaFold3 (AF3) modelling and X-ray crystallography, we identify two collagen-like proteins that are involved in F-type fiber biogenesis. We elucidate the assembly mechanism by unravelling the molecular mode of docking to the exosporium and the mode of fibrillum decoration. Phylogenetic analysis demonstrates that F-ENA is widespread in the class of Bacilli and is to a lesser extent also found in the class of Clostridia. Moreover, we identify a second subgroup of ruffle-less F-like ENA’s in the *Paenibacillus* genus and solve the cryoEM structure of a representative member.

## Results

### *Bacillus thuringiensis* subsp kurstaki spores are decorated with F- and S-type endospore appendages

To gain insight into the molecular architecture of a Btk spore biofilms, a confluent macrocolony of sporulated Btk was investigated in negative stain transmission electron microscopy (nsTEM). Btk spores were observed to be decorated with two types of endospore appendage (ENA) types: S-ENA, with a 10 nm diameter and typical staggered 2D projection image (Supporting Figure 1, 3), recently described for *B. paranthracis*^14^ and a second, unidentified ENA measuring 5 nm in diameter (Figure 1, Supporting Figure 1). Given the increased flexibility of these unknown ENAs compared to the more rigid S- and L-type ENAs^14,16^, we hitherto refer to them as ‘fibrillar <u>e</u>ndospore <u>a</u>ppendages’ or in short F-ENA. Btk spores are enclosed by an exosporium. High-magnification imaging of the hairy nap (BclA layer) shows that F-ENAs are anchored to the exosporium base layer and protrude through the hairy nap (Figure 1b). At the spore’s distal terminus, F-ENAs are decorated with a 3 nm diameter flexible tip fibrillum or ‘ruffle’, which is characterized by a 200 nm long stalk region ending into a distinctive globular tip or head-group (Figure 1c). We further demonstrate that another widely used Bt strain, i.e. Bt subsp. Berliner (Btb), is also abundantly decorated with F-ENAs that terminate into a tip fibrillum (Figure 1d). Interestingly, the F-ENA ruffles of Btb are shorter than the Btk F-type ruffles, measuring 85 nm in length. The presence of ruffles and the observed differences in length are reminiscent to the termini of S- and L-ENAs found on *B. paranthracis*, which terminate into multiple ruffles (e.g. 3-4) or a singular ruffle, respectively^14,16^. The presence of ruffles on the S-ENA fibers of Btk is also confirmed in this study. Using nsTEM, we resolved multiple 125±6nm (n=12; mean; standard deviaton; 1-3 ruffles per S-ENA fiber) long ruffles on Btk S-ENA (Supporting Figure 1).

**Figure 1.**
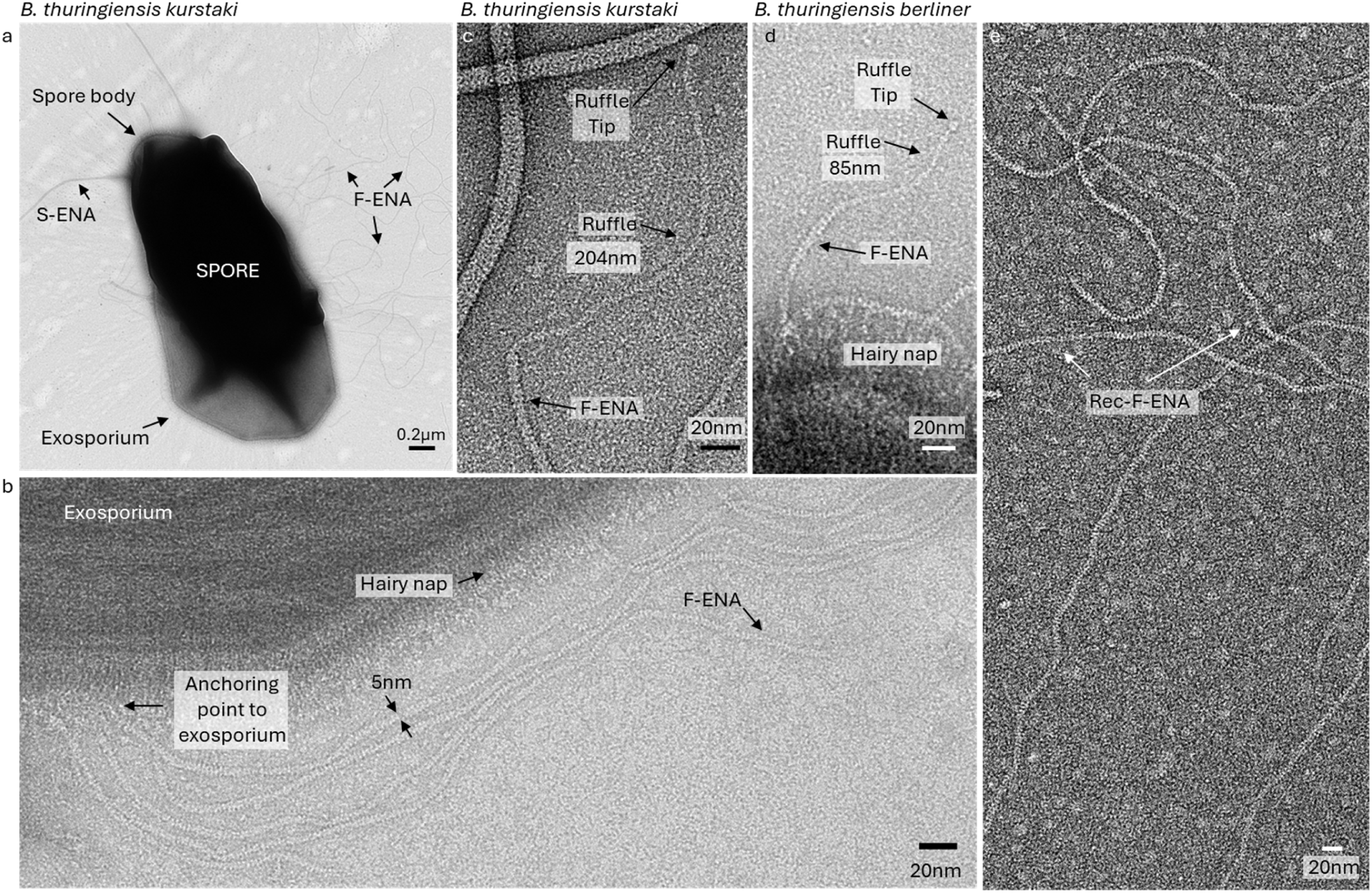
nsTEM observations of Bt kurstaki and berliner spores revealing F-ENA fibers. (a) Low magnification micrograph of *Bacillus thuringiensis* subspecies kurstaki decorated with previously identified S- and newly discovered F-type endospore appendages (F-ENA). (b) High magnification micrograph of F-ENA (diameter = 5 nm) anchored to the exosporium, permeating through the hairy nap layer. (c,d) The spore distal F-ENA terminus is decorated with a flexible 3nm diameter tip-fibrillum (ruffle) that terminates into a globular head group. For Btk (c) the ruffle measures ±200 nm, whereas for Btb (d), the ruffle is 85 nm long. (e) micrograph of recombinantly produced F-ENA fibers purified from the cytoplasm of *E. coli*.

### CryoEM structure of F-ENA and S-ENA of *Bacillus thuringiensis* subsp kurstaki

Next, we aimed to characterize F-ENAs using cryogenic transmission electron microscopy (cryoEM). To achieve this, we developed a protocol to shear off the F-ENA fibers from the spore surface. The resulting ENA suspension was plunge-frozen and a cryoEM dataset was acquired using a cryoARM 300 microscope (Figure 2b). Using standard helical processing in cryoSPARC, we obtained reconstructed cryoEM volumes with resolutions of 2.7 Å (based on the FSC 0.143 criterion) and 4.1 Å for F- and S-ENA, respectively (Figure 2a,i; Supporting Figure 2). Helical parameters for F-ENA were estimated from the 2D class averages (Figure 2b), while the rise and twist values of 7A02 (PDB) were used as inputs for S-ENA. The refined helical parameters were determined to be a twist of 42.1° and a rise of 33.7 Å for F-Ena, and a twist of −31.1° and rise of 3.14 Å for S-ENA.

**Figure 2.**
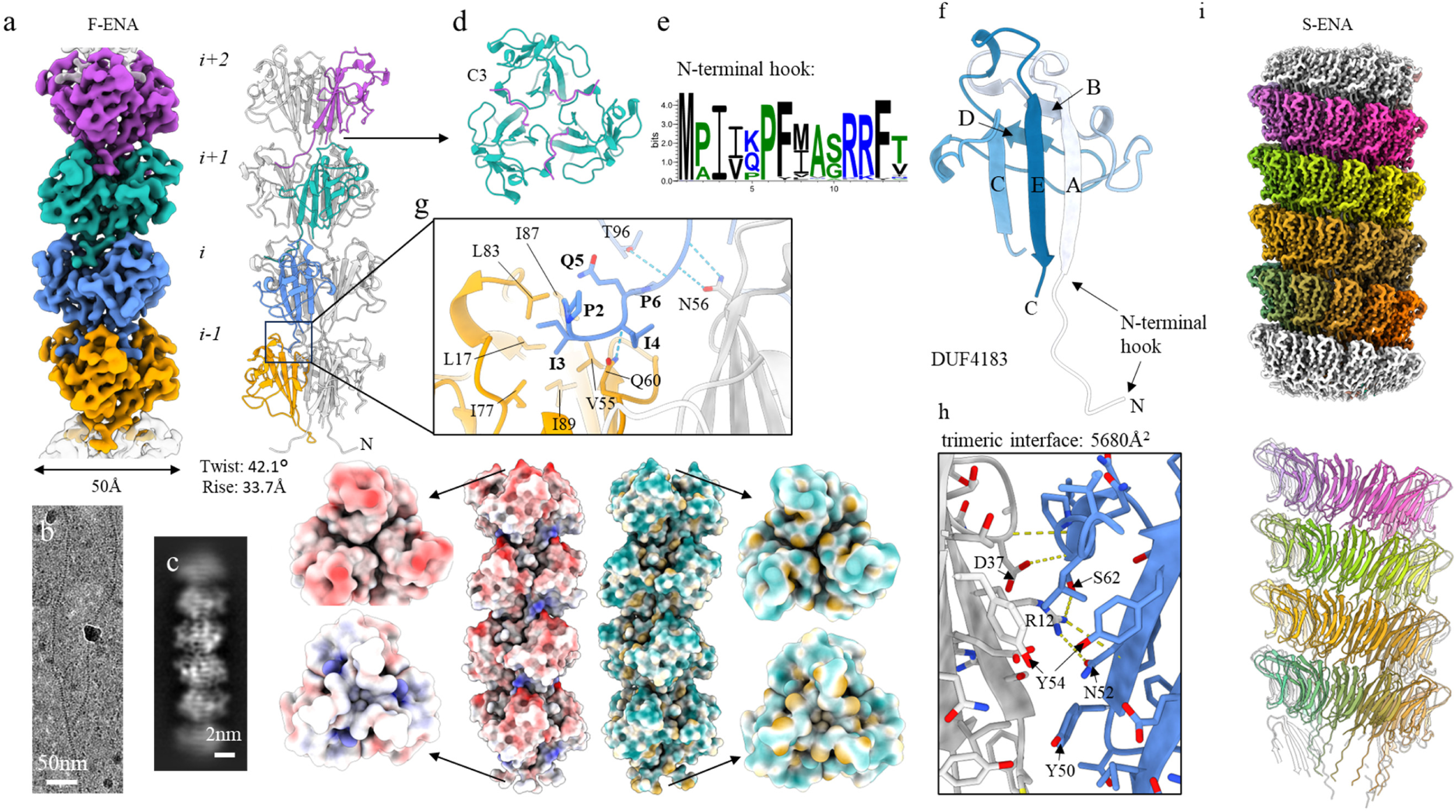
CryoEM analysis of F-ENA fibers found on the surface of *Bacillus thuringiensis* kurstaki spores. (a) Reconstructed cryoEM volume (colour coded per fiber segment which corresponds to an F-ENA trimer) and corresponding molecular model of F-ENA. (b) 60k magnification cryoEM micrograph of an F-ENA fiber purified from Btk. (c) 2D class average of extracted F-ENA fiber segments. (d) Top-view image of the F-ENA pilus. (e) Cartoon representation of a single F-ENA protomer which belongs to the DUF 4183 family characterized by two juxtaposed beta-sheets interconnected by a series of loops, and an N-terminal extension (Ntc) that serves as the docking mechanism for incoming F-ENA trimers to connect to the fiber terminus. (f) Zoom-in of the Ntc of segment *i* docked into the receiving hydrophobic groove of segment *i-1*.

Inspection of the F-ENA volume revealed a trimeric symmetry, and thus processing was continued by applying C3 symmetry along the Z-axis (Figure 2a). The resulting volume displayed clear, continuous density of the protein backbone, but did not permit precise de novo assignment of all sidechain entities. To facilitate unambiguous protein identification, the purified fiber suspension was subjected to trypsin digestion, followed by liquid chromatography tandem mass spectrometry (LC-MS-MS). The initial list of 276 detected proteins was filtered according to protein size. Using a *de novo*-built main chain model generated by Modelangelo^17^, we estimated that the major F-ENA subunit would be approximately 100 amino acids (aa) in length, and hence only proteins detected in the LC-MS-MS analysis with lengths between 70 and 130 aa were retained for further analysis. From the 38 retained sequences, we retrieved the corresponding AlphaFold2 models from the AlphaFold Protein Structure Database. Pairwise structural alignments, performed using FoldSeek with a *de novo* F-ENA model as a template, led to the unambiguous identification of WP_001121647.1 as the major subunit of F-type fibers in Btk. WP_001121647.1, hereafter called F-ENA, is a 96 aa protein that is classified by Pfam as a member of the Domain of Unknown function 4183 (DUF4183; PF13799). To confirm the ability of WP_001121647.1 to self-assemble into F-ENA fibers, we recombinantly expressed the protein in the cytoplasm of *E. coli*. nsTEM analysis of the insoluble fraction of the lysate revealed the presence of 5 nm diameter flexible fibrils that were indistinguishable from *ex vivo* F-ENA fibers (Figure 1e).

The Btk F-ENA fold comprises two juxtaposed β-sheets composed of strands A, C, E and B, D respectively, along with a short N-terminal extension (Figure 2f). Using the WP_001121647.1 sequence as input, an atomic model of the helical architecture of F-type fibers was constructed **Figure 2** using Modelangelo, followed by real-space refinement in Coot and Phenix (**Figure 2**a). F-ENA subunits assemble into trimeric units that stack axially with a 42° rotation (Fig.2a,d). Lateral stacking at the trimeric interface occurs through a predominantly polar interaction with an interface area of 5680Å^2^, involving 7 identified hydrogen bonds, mainly mediated by sidechains (Figure 2h). Axial stacking is principally facilitated by the docking of an N-terminal 7 residue extension, referred to as the ‘N-terminal hook’, into a cleft defined by the interface between two neighbouring F-ENA protomers in the trimer configuration (Figure 2e-g). The interaction between the N-terminal hook of subunit *i* and protomers of the preceding fiber segment *i-1* is mainly composed of hydrophobic contacts, where residues P2, I3 and I4 dock into a hydrophobic groove lined by small hydrophobic sidechains, including L17, V55, I77, L83, I87 and I89 (Figure 2g). These N-terminal hook residues are conserved, as judged from the consensus logo derived from a multiple sequence alignment of the top 500 blast hits using WP_001121647.1 as the query (Figure 2e). Guided by the cryoEM structure, we identify the following signature motif for the F-ENA N-terminal hook region: M-P-Ψ_1_-Ψ_2_-$-P-Ψ_3_ where Ψ_1_ and Ψ_2_ represent small hydrophobic residues (I,V), Ψ_3_ tends to be a larger hydrophobic, aromatic residue (F,Y) and $ is a surface-exposed polar residue (K,Q). Examination of the surface electrostatics of the F-ENA trimer (https://server.poissonboltzmann.org/) reveals a net negative charge of −6 at pH 7.0 (PDB2PQR). Notably, residues D81 and D84 form a negative patch on the surface of the spore distal terminus, repeated threefold due to the C3 symmetry (Supporting Figure 4). In contrast, at the spore proximal F-ENA terminus, a surface-exposed arginine (R11) forms a salt-bridge with D84 of the preceding segment *i-1* (Supporting Figure 4d). Using the dipole moment server (https://dipole.proteopedia.org/), we calculated an overall molecular dipole moment for the F-ENA trimer of 772 ± 0.55 D (Supporting Figure 4b), which is aligned with the helical axis (Supporting Figure 4c). This suggests that new F-ENA trimers that are diffusing towards the fiber tip may become pre-oriented via electrostatic interactions (driven by the alignment of the dipole moment) and subsequently latch onto the fiber tip via coordinated docking of the three N-terminal hook segments into the corresponding hydrophobic clefts. Despite the relatively small interaction interface, the F-ENA fibers are relatively stable. Testament to that stability is the fact that the fibers were washed numerous times in miliQ water prior to cryoEM analysis, showing no clear signs of depolymerization.

The limited resolution (4.1 Å) of the S-ENA cryoEM volume prevented us from unambiguously identifying the constituent subunits of the S-ENA Btk fiber (Figure 2i). Recently, we showed that *ex vivo* purified S-ENA from *Bacillus paranthracis* are composed of two different subunits, Ena1A and Ena1B. Considering that the relative stoichiometry of Ena1A and Ena1B (which share 39.4% sequence identity) was unknown, and both subunits are likely randomly distributed along the length of the fiber, the reconstructed EM-volume represents a weighted average of both sequences, stopping us from building an atomic model for the *ex vivo* S-type fibers. To address this issue, we solved the structure of recombinant Ena1B fibers. Anticipating similar ambiguities during the model building for *ex vivo* Btk S-ENA, we produced recombinant homo-polymeric S-ENA by expressing WP_001277547.1 (hitherto referred to as Ena2A), which shares 37.8% sequence identity with the solved Ena1B structure (PDB:7A02), in the cytoplasm of *E. coli*. After purifying S-ENA from the *E. coli* lysate, we collected a cryoEM dataset and solved the Ena2A structure to 2.7 Å resolution (Figure 2i; Supporting Figure 3). The helical parameters for Ena2A were refined to a twist of 31.03° and a rise of 3.17 Å, which is subtly different to the rise and twist of Ena1B, i.e. −32.35° and 3.44 Å respectively (Supporting Figure 5). At the protomer level, the pruned and unpruned atomic root mean squared deviations (RMSD) between Ena2A and Ena1B are 0.9 Å and 2.6 Å, with the most significant structural differences located in the N-terminal connector (residues 8-15). The positioning of the double cysteine motif (C_12_C_13_) is slightly closer to the Ena2A fold compared to Ena1B. This is the result of the shorter linker region to strand B (Ena2A: P_14_D_15_ / Ena1B: A_14_N_15_G_16_Q_17_), which translates into the observed reduction in rise. Similarly to Ena1B, we could not resolve the first 8 residues, suggesting that this region is flexible or disordered; and its role in fiber biogenesis remains unclear. Despite the low sequence identity, there is remarkable structural conservation between the two structures. We attribute this structural preservation to the mode of protomer coupling, i.e. β-sheet augmentation. Pairwise docking of open-ended sheets entails the formation of a hydrogen bonding network between the main chains of both protomers and can therefore reasonably be expected to be insensitive to the protein sequence, provided the main fold and overall surface characteristics (such as charge and hydrophilicity) are conserved.

### Two collagen-like proteins are required for F-ENA coupling to the exosporium and for the formation of the tip fibrillum

Inspection of the local genomic context of the *f-ena* gene (locus: BTK_RS13480 in NZ_CP004870) showed that the upstream neighbouring gene (BTK_RS13475) is predicted to encode a 327-residue protein (M1Q906), hitherto referred to as F-anchor (nomenclature explained below; Figure 3a). An InterProScan^18^ analysis of the primary sequence identified three distinct domains: an N-terminal exosporium leader peptide (residues 9-37), a collagen triple helix repeat region (residues 30-241, comprising 70 GXY triplet repeats) and a C-terminal DUF4183 domain (residues 252-323). An exosporium leader sequence was first identified in Bcl^19^ and has been shown to be responsible for the tethering of the BclA N-terminus to the exosporium via docking to ExsF, forming the hairy nap^20^. The presence of an exosporium leader sequence on the N-terminus of F-Anchor fits with the experimentally observed localization of F-ENA on the surface of the exosporium (Figure 1b).

**Figure 3.**
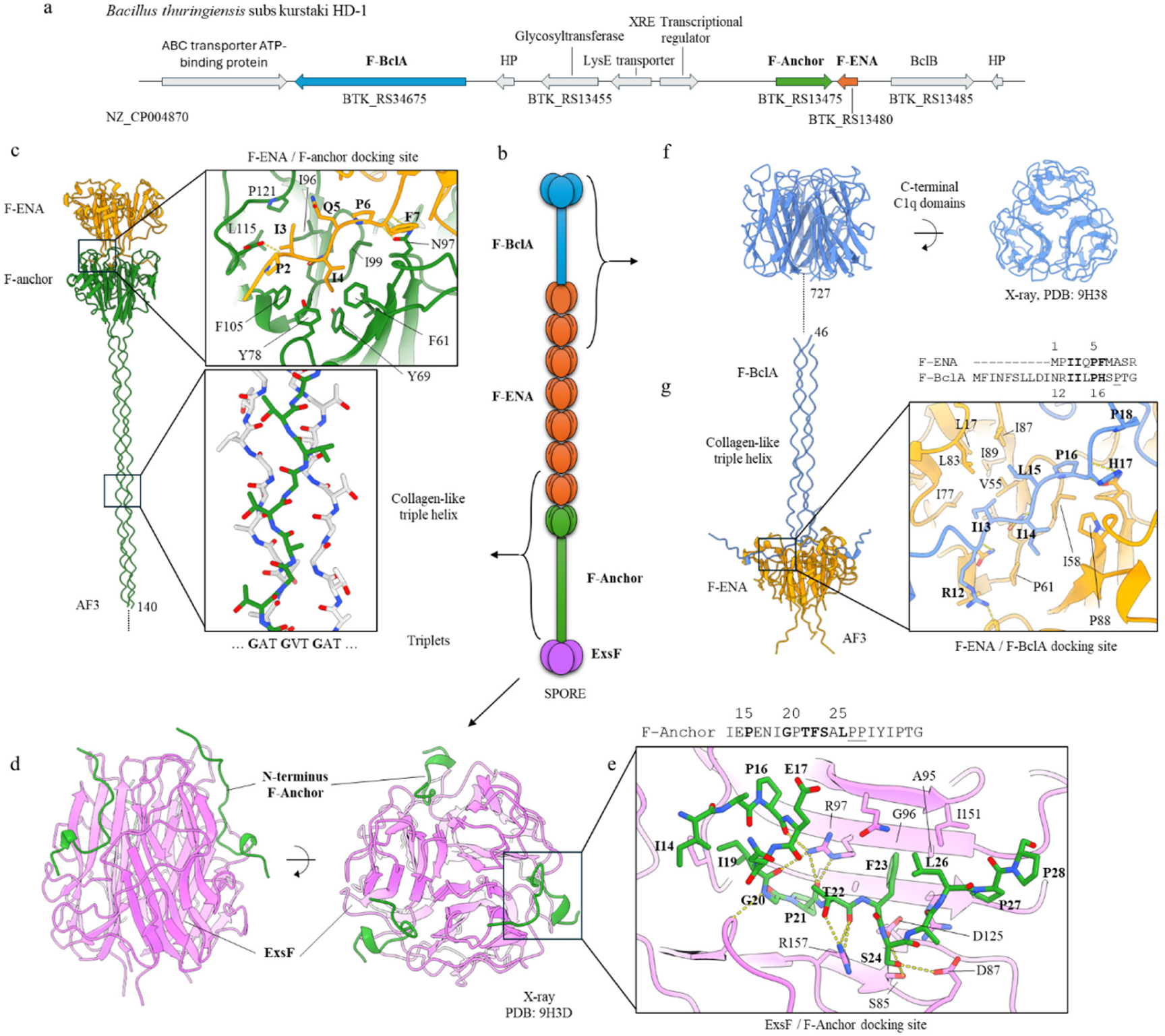
Structural organization of the F-type endospore appendages. (a) Genetic locus of the genes associated with the biogenesis of F-ENA; (b) Schematic representation of the F-type pili tethered at the spore-proximal end to the exosporium via a tripartite F-ENA / F-Anchor / ExsF complex, and decorated at the spore-distal terminus with the tip-fibrillum F-BclA; (c) AlphaFold3 prediction of the F-ENA / F-Anchor complex. A truncated F-Anchor containing residues 140 to 276 was used for clarity purposes; (d) X-ray crystal structure of ExsF in complex with a peptide (F-Anchor_14-28_) corresponding to residues 14 to 28 of F-Anchor; (e) Zoom-in of the ExsF / F-Anchor_14-28_ interaction; (f) X-ray crystal structure of the C-terminal head group (F-BclA_727-863_) of F-BclA; (g) AlphaFold3 prediction of the F-ENA / F-BclA_1-46_ complex.

To test whether the putative exosporium leader sequence of F-Anchor would indeed form a complex with ExsF, we co-crystallized recombinantly produced and purified ExsF, which was tagged with an N-terminal 6xHis tag, with a synthetically produced peptide spanning residues 14 to 35 of F-Anchor (NH_2_-IEPENIGPTFSALPPIYIPTG-COOH; annotated as FA_14-35_) (Supporting Figure 6,7). The crystal structure of the ExsF/FA_14-35_ complex was solved to 1.92 Å resolution (R/R_free_ of 18.4/21.6) and reveals three FA_14-35_ copies that are docked into a cleft at the trimeric interface between two neighbouring ExsF copies (Figure 3f; Supporting Figure 7a). The interaction between FA_14-35_ and ExsF involves both polar and apolar contacts, with no salt bridges detected. Three ExsF interacting regions can be identified in FA_14-35_ (Supporting Figure 7a and 7b). In region 1, residues I14, P16 and I19 tether the N-terminus of the peptide to a hydrophobic patch on the surface of ExsF (Supporting Figure 7a). Within region 2, we identified 14 hydrogen bonds between FA_14-35_ and ExsF, involving both sidechain-sidechain and sidechain-mainchain contacts. Notably P16, I19 and P21 are hydrogen bonded to R97, T22 interacts with R157, and S24 forms H-bonds with D87 and S85 of ExsF. In addition, F23 fits tightly into a hydrophobic pocket lined by I151, the side-chain of Q149 (Cγ), A95, and the Cβ-Cδ region of R97. Towards the C-terminal end of FA_14-35_ (region 3), residues L26 and I29 make hydrophobic contacts with I15, P93 and V91, while Y30 forms multiple hydrogen bonds with N119 and A92. Although the N-terminal sequence of F-Anchor is strongly conserved (Supporting Figure 7b; Supporting Figure 8a), the role of the first 13 residues remains unclear. For BclA, it has been shown that residues 1-19are proteolytically cleaved off^19^ and do not contribute to binding to ExsF. The pairwise sequence alignment between the N-termini of BclA and F-Anchor shows clear sequence similarity (Supporting Figure 7c), suggesting that they may share similar modes of binding to ExsF. Whether residues 1 to 13 from F-Anchor also get proteolytically removed is not known. We also looked at the consensus motif for the general exosporium leader peptide, which we derived from the hmm profile of NCBifam entry TIGR03720 (Supporting Figure 7d). Conservation in the motif is highest at positions 12 to 17. These positions map onto the region that forms the greatest number of H-bonds with ExsF. We also note strong conservation of the proline residues at positions 9, 14, 17, 18, 20 and 21. Based on the ExsF/FA_14-35_ crystal structure we can rationalize this conservation. These positions correspond to kinks in the peptide backbone that are required to accommodate the meandering mainchain path imposed by the shape of the cleft defined by the trimeric ExsF interface.

F-Anchor shares many similarities with BclA. Both are collagen-like proteins and both couple to the exosporium of Bt. The key distinction, however, lies in their respective C-terminal domains. BclA has a ‘BclA-C domain’ (Pfam: PF18573; TNF/C1q superfamily: SSF49842), whereas F-Anchor possesses a C-terminal DUF4183 domain. This divergence suggests that F-Anchor will likely serve a different function compared to BclA. Although the pairwise sequence identity of the DUF4183 domain of F-anchor to F-ENA is only 23%, we reasoned that F-anchor and F-ENA might form a hetero-hexameric complex. AlphaFold3^21^ (AF3) predictions support this hypothesis, showing a high-confidence dimer of trimers (3 F-ENA / 3 F-anchor: ipTM = 0.89, pTM = 0.91) that mimics the homotypic interactions between F-ENA ring segments in the F-type pilus (Figure 3b). In the AF3 model, the N-terminal hook of F-ENA docks into a solvent-exposed hydrophobic cleft on the surface of the F-anchor trimer. The pruned RMSD between the DUF4183 domains of the F-ENA trimer and the F-anchor trimer is 0.8 Å, indicating a high degree of structural similarity. When considering all atom pairs (non-pruned), the RMSD increases to 3.5Å, reflecting minor variations likely due to differences in sidechain conformations and flexibility. This close structural agreement supports the potential for hetero-hexameric complex formation.

These data strongly suggest that F-Anchor functions as a critical bridging moiety that tethers F-ENA to the exosporium of Btk by displaying an F-ENA like C-terminal domain (CTD) on a semi-flexible display platform (i.e. the collagen stalk). We hypothesize that F-Anchor also serves as a starting point for F-ENA self-assembly, in that the F-Anchor-CTD will capture incoming F-ENA trimers through conserved docking, to preferentially initiate fiber formation onto the exosporium. To ensure efficient F-ENA capture, F-Anchor must remain sterically accessible despite being integrated in the hairy nap. The collagen region of BclA spans residue 29 to 161, corresponding to 40 triplet repeats. Based on AF3 predictions, the expected length of a collagen-helix segment is approximately 2.8 Å/residue. For F-Anchor and BclA this translates into a collagen-helix length of 59 nm and 37 nm, respectively. Using nsTEM, we measured the width of the Btk hairy nap layer to be 30 nm, which is in correspondence with the expected dimensions of BclA, assuming full extension of the collagen stalk. F-Anchor can therefore be expected to protrude through the hairy nap, making it fully accessible for binding with F-ENA. Sylvestre *et al* showed that polymorphism in the collagen-like region of BclA leads to variation in the width of the hairy nap layer^13^. This raises the question of whether similar polymorphism in the collagen region of BclA could be coupled to potential polymorphism in F-Anchor. Hence, to test whether the observed difference in length between F-Anchor and BclA of Btk is a conserved feature for all F-ENA harbouring strains, we downloaded all the NCBI bacterial genomes in the Bacillus/Clostridium group (taxid 1239, assembly level complete) and used hmmersearch (E-value < 1E-40) to identify F-ENA and BclA homologue pairs. In Supporting Figure 9 we plot the lengths of the respective collagen stalks (expressed in number of amino acids) which shows that the stalk region of F-Anchor is, on average, longer than that of BclA. This suggests that the collagen stalk polymorphism of both proteins could indeed be coupled, with a bias toward CLS regions that are longer in F-Anchor than in BclAs.

In addition to F-Anchor, we identified another gene possibly involved in F-ENA biogenesis: BTK_RS34675. This gene also encodes a putative collagen-like protein with C-terminal C1q domain (Figure 3a) and is referred to as F-BclA (see below). Interproscan does not identify an N-terminal exosporium leader sequence in F-BclA, indicating it will likely not be coupled to the exosporium. A pairwise-sequence alignment of the F-BclA N-terminus (F-NTD) with the F-ENA N-terminal hook sequence reveals a putative F-ENA docking motif in F-BclA (starting at residue 14: IILPH) that follows the Ψ_1_-Ψ_2_-$-P-F pattern (observed in F-ENA as IIQPF) responsible for F-ENA/F-ENA docking (Figure 3g). Based on this analysis, we reasoned that F-BclA might encode for the tip-fibrillum of F-ENA. Indeed, AF3 predicts a heterohexameric F-ENA/F-BclA complex that mimics the homotypic F-ENA interactions. Specifically, the putative F-BclA docking motif (IILPH) is modeled by AF3 to bind at the same location as the F-ENA N-terminal hook. Moreover, the collagen region of F-BclA comprises 701 residues, which translates into an extended triple-collagen helix of 196 nm in length. This fits with the experimentally measured value of 204 nm for the F-ENA of Btk. Next, we recombinantly expressed, purified and crystallized the C-terminal domain of F-BclA (F-BCLA-CTD; Supporting Figure 10). Using X-ray crystallography, we solved the structure of F-BCLA-CTD to 2.46Å resolution (R/R_free_ = 22.5/27.7). F-BCLA-CTD forms a trimeric complex of all-β domains with a TNF-like jelly fold topology, reminiscent of the C-terminal domain of BclA (BclA-C, PDB entry 1WCK). The RMSD between F-BCLA-CTD and BclA-C is 0.7Å, with a sequence identity of 27.5%.

BclAs from *B. cereus* and *B. anthracis* have been shown to be extensively O-linked glycosylated in a domain specific manner^22^. In these proteins, the collagen regions are modified with species specific short O-glycans (anthrose, cereose) and the BclA-C domains are substituted by species specific polysaccharide O-linked glycans. Although the F-BCLA-CTD presented here is not glycosylated as a result of recombinant expression in *E. coli*, we anticipate that F-BclA likely will be glycosylated in its native context. Firstly, two genes upstream of F-BclA lies the gene *BTK_RS13455,* which encodes for a putative glycosyltransferase (Figure 3a). Secondly, F-BCLA-CTD contains a remarkable 78 out of 185 (42%) solvent exposed residues that are serine or threonine. Finally, the collagen region of F-BclA (residues 19 to 720) consists of 326 threonine or serine residues, accounting for 45% of the collagen stalk (Supporting Figure 11). We also analysed the conservation of F-BclA by performing BLAST searches using either F-NTD or F-BCLA-CTD as query sequences. Both approaches lead to very similar results, producing a dataset of collagen-like sequences in which both the N- and C-terminal domains are highly conserved (see Supporting Figure 8). Note that searches using F-NTD as a query could have identified putative F-ENA ruffle genes with a different C-terminal domain, but no such sequences were detected using this approach.

Taken altogether, a picture emerges of F-ENA fibers that are coupled to the exosporium via a dedicated anchoring complex (ExsF/F-Anchor/F-ENA) on the spore proximal side, and decorated with a dedicated, conserved collagen-like (putatively glycosylated) tip fibrillum (F-BclA) on the spore distal side that extends into the surrounding environment (Figure 3b).

### F-ENA contributes to spore clustering

We recently showed that S- and L-type endospore appendages in *B*. *paranthracis* contribute to spore-spore interactions that lead to the formation of spore clusters^23^. In this study, we tested if the endospore appendages of *B*. *thuringiensis* are also involved in the auto-aggregation of spores. nsTEM micrographs of freshly prepared Btk spore solutions deposited onto a Cu-mesh grid revealed the presence of loosely packed spore clusters wherein spores typically do not engage in direct lateral contacts via the spore body or the exosporium (Figure 4a). Higher magnification imaging revealed a dense network of F- and S-ENA-fibers pervading the spaces in between adjacent spores (Figure 4b,c). It is tempting to speculate at this point that ENA is mediating direct spore contacts by binding to a (specific) surface receptor on neighbouring spores. However, it is important to stress that the observed globular spore clusters could also have formed during the EM grid preparation process (i.e. as a drying artefact), and may not accurately reflect the spore assembly state in solution. In addition to globular spore clusters, however, we also observed linear trains of spores spaced at semi-regular intervals (seemingly) connected by S-type ENAs (Figure 4d). Further high-magnification imaging also revealed F-ENA fibers bridging the interstitial regions between neighbouring spores (Figure 4e). We argue that such linear spore trains, which are in an extended state, likely formed as a result of viscous drag during the sample blotting process and are indeed indicative of inter-spore clustering and contacts in the liquid phase.

**Figure 4.**
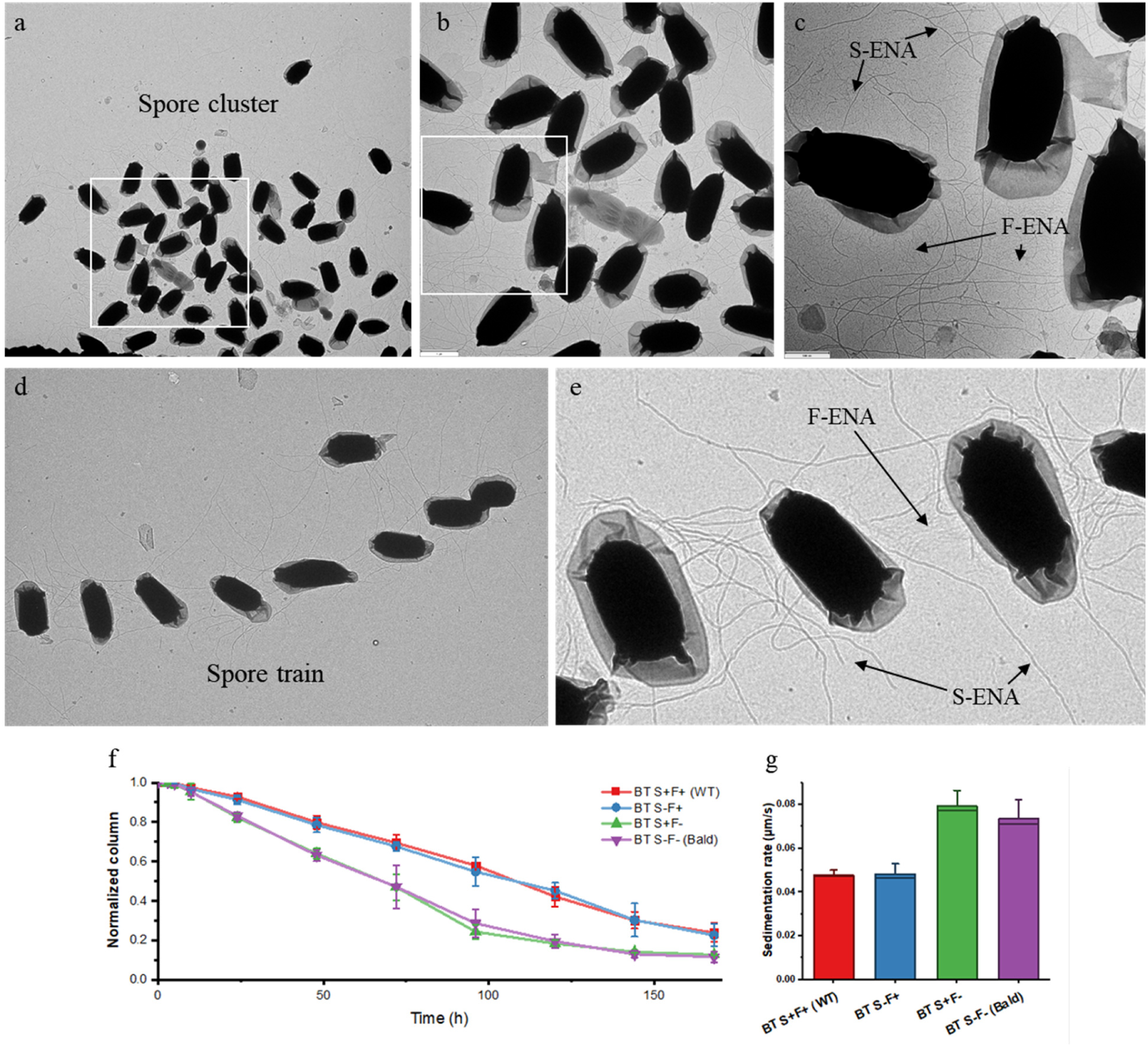
ENA-mediated aggregation of Btk spores. (a)-(c) nsTEM image of a globular Btk spore cluster at consecutively higher magnification (white rectangle denotes zoom-in area). F- and S-type ENAs populate the regions in between the spores; (d) Linear spore train of Btk spores; (e) S- and F-type fibers making lateral contacts across neighbouring spores; (f) Time dependence of the location of the developing sedimentation front; (g) Mean rate of sedimentation (µm/s).

To probe for the relative contributions of S- and F-type ENAs in spore clustering, we set out to generate deletion strains deficient in either S-type (Bt S^-^F^+^), F-type (Bt S^+^F^-^) or both ENA types (Bt S^-^F^-^). However, we experienced difficulties in producing ENA-knockouts in Btk, and had to resort to knockout production in an alternative strain, i.e. *B. thuringiensis* 407 (taxid: 527021; ASM16149v1). The Bt407 strain is closely related to Btk and produces F- and S-ENA fibers with high similarity (Ena1A: 76.99%, Ena1B: 85.37%, and F-ENA: 100% sequence identity). All Bt407 deletion strains and the WT Bt407 strain were sporulated on solid media (LB agar) through nutrient starvation, harvested, resuspended in Mili-Q water and stored under stagnant conditions. Time-lapse photography was used to monitor the sedimentation kinetics over a 175h period by tracking the position of the interface that develops between the spore dense and dilute region (Figure 4f). We see a marked increase in the average rate of sedimentation for Bt S^+^F^-^ compared to WT and Bt S^-^F^+^. This observation would be in agreement with F-ENA ‘agglutinating’ suspended spores into a loosely contacting network. Notably, we did not record an additional increase in sedimentation rate for the double knockout (S^-^F^-^) compared to Bt S^+^F^-^, suggesting F-ENA takes a dominant role in the agglutinating activity of the Bt407 extrasporal matrix.

### Paenibacillacea spores are decorated with a subclass of F-ENA-like pili

To understand the phylogenetic distribution of F-ENA, we searched for DUF4183-containing genes across the *B. cereus* group sensu lato using the btyper database. Out of the 5976 genomes searched, 5647 (94.4%) harbour one or more DUF4183-containing genes (Supporting Figure 12a). Looking more broadly across the Bacillota phylum (taxid: 1239), we examined 9342 complete Genbank genomes, of which 783 (8.3%) hold one or more DUF4183-containing genes (Supporting Figure 12b). These F-ENA carrying genomes cluster into the Bacillacea family, containing the *B. cereus* group hits, *Lysinibacillus*, *Geobacillus*, *Priestia*, and *Anoxybacillus*, as well as the Paenibacillaceae family, which includes hits in *Paenibacillus*, *Brevibacillus* and *Cohnella*. A limited number of hits were also obtained in the Clostridia class.

Looking at the Paenibacillaceae hits in further detail, we identified F-ENA homologues with N- and C-terminal extensions to the DUF4183 domain. A0A1I1C8X4 (22.7% sequence identity to F-ENA; hereafter referred to as F-ENA-2a; 136aa) is one such example, for which AF3 predicts extensions to the A and E β-strands (Supporting Figure 12b). To test whether this distant homologue still self-assembles into F-ENA-like fibers, we recombinantly expressed and purified F-ENA-2a in the cytoplasm of *E. coli*. nsTEM micrographs of the insoluble *E. coli* fraction after cell lysis confirmed the presence of 3 nm diameter fibrils with a regularly spaced ‘pearl on strings’ appearance, and which exist either as single fibers (Figure 5a) or are grouped into parallel bundles (Figure 5c,d). We collected a cryoEM dataset and solved the F-ENA-2a pilus structure to a global resolution of 4.1Å, with a helical rise and twist of 95.25Å and 67.50°, respectively (Figure 5e). Due to the limited resolution, we sharpened the reconstructed volume with EMready^24^ and performed rigid-body docking using an AF3 F-ENA-2a trimer as the starting model (Figure 4f). Similar to F-ENA, F-ENA-2a fibers are helical ultrastructures with global C3 symmetry, where each fiber segment (asymmetric unit) consists of an F-ENA-2a trimer that couples to the next segment via docking of the N-terminal hooks (Figure 5i,g). Like F-ENA, the F-ENA-2a N-terminal hook is composed of the first 7 residues (<u>MP</u>**VI***K*<u>P</u>**V**…) in a conserved <u>M</u>-<u>P</u>-**Ψ_1_**-**Ψ_2_**-*$*-<u>P</u>-**Ψ_3_** motif, that adheres to the hydrophobic groove on the head domain of a preceding segment. The latter is composed of a globular DUF4183 region that is structurally equivalent to the DUF4183 fold of F-ENA (pruned RMSD = 1.0 Å). The N- and C-terminal extensions of F-ENA-2a form a toroidal β-barrel extension or ‘neck’ composed of 6 antiparallel strands that position the N-terminal hooks 110 Å away from the head domains. This configuration results in the characteristic dotted pearl on strings appearance observed in cryoEM and nsTEM micrographs (Figure 5a-d,h). Despite the low sequence identity to F-ENA, F-ENA-2a is characterized by a similar bipolar distribution of surface charges (Figure 5j), leading to a dipole moment of 2561D, with the negative pole situated at the head group and directed towards the positive N-terminal hook region (Supporting Figure 14). Similarly to F-ENA, three salt bridges reinforce inter-segment connections between the head groups of segment *i* and the N-terminal hooks of segment *i+*1, specifically between K5 and D101.

**Figure 5.**
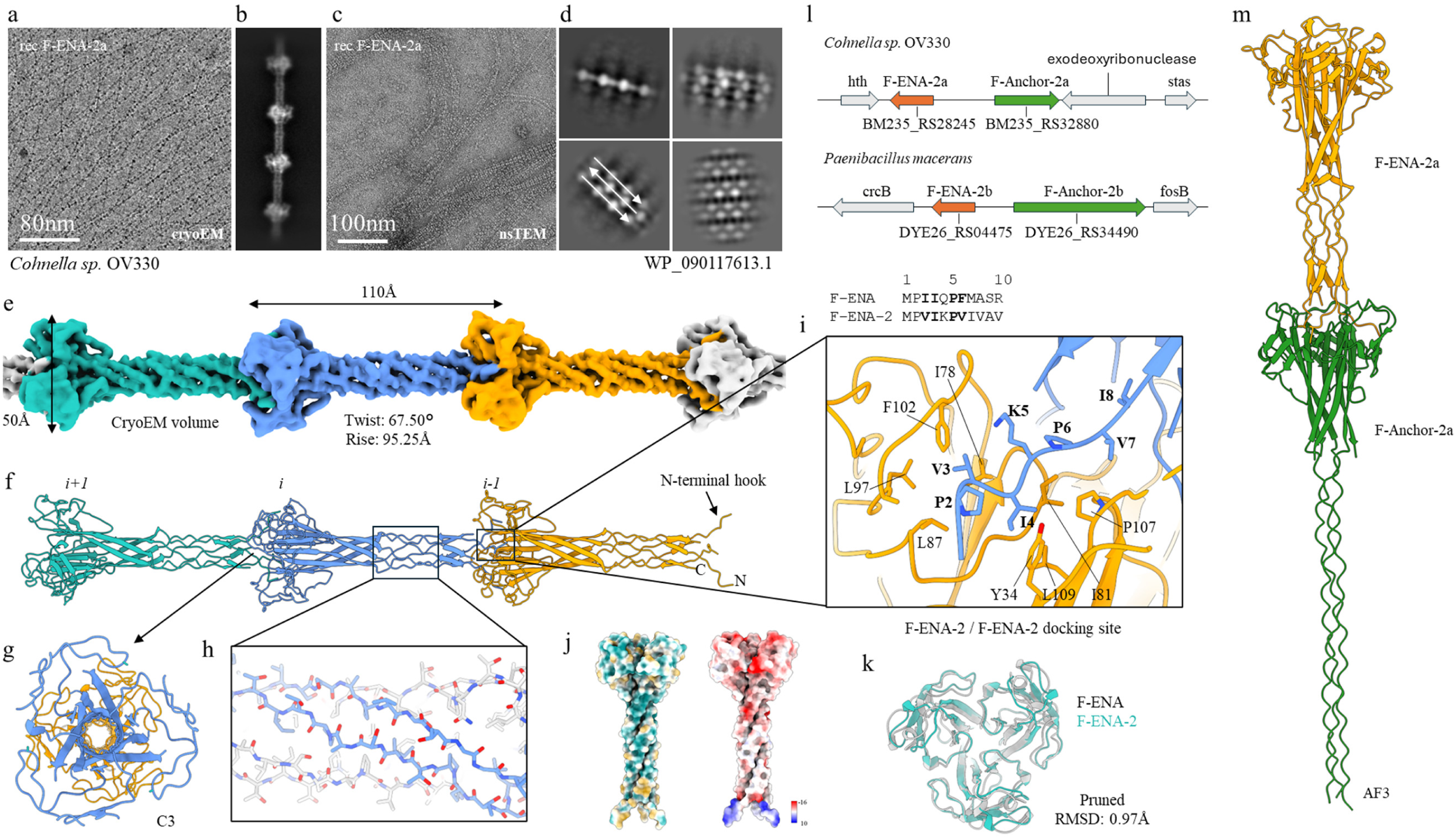
CryoEM of F-type-2 ENAa fibers found on the surface of Paenibacillaceae spores. (a) cryoEM micrograph of rec-F-ENA-2a with corresponding 2D class average in (b); (c) nsTEM micrograph of rec-F-ENA-2a with corresponding 2D class averages in (d); (e) reconstructed cryoEM volume with (f) the corresponding molecular model shown in cartoon representation; (g) top-view slice-through along the C3 axis; (h) zoom-in of the beta-cylinder extension; (i) docking site of the N-terminal hook; (j) hydrophobic and electrostatics surface colouring; (k) structural overlay of the DUF4183 domains of F-ENA and F-ENA-2a; (l) genetic loci of F-ENA-2 for two members of the Paenabillacaea group; (m) AF3 model of the F-ENA-2a and F-Anchor-2a complex

Looking at the genomic context of *f-ena-2a* (Refseq locus BM235_RS28245 in NZ_FOKE01000015.1 of *Chonella* sp. OV330), we identified the gene BM235_RS32880, which encodes WP_218161362.1, a putative DUF4183-domain containing protein (Figure 4l). Inspection of the primary sequence reveals that this protein contains collagen-repeats. AF3 predicts a putative hetero-hexameric complex between F-ENA-2a and WP_218161362.1 that is similar to the predicted complex of F-ENA and F-Anchor (Figure 5m). Based on this observation, we hypothesize that WP_218161362.1 may serve a similar role in coupling the F-ENA-2a pili to the spore, and for this reason we refer to WP_218161362.1 as F-Anchor-2a (23% sequence identity to F-Anchor). We did not identify a putative ruffle for F-ENA-2a, suggesting these fibers lack a tip fibrillum.

It is interesting to note that recombinant F-ENA-2a fibers can form intercalated super bundles, wherein the fibers stack laterally in an anti-parallel fashion (Figure 5c,d, Supporting Figure 15). To test whether the F-ENA-2a fiber isoform can also be found in an *in vivo* spore setting, we screened various *Paenibacillus* strains and observed F-ENA-2a-like fibers on one of the *Paenibacillus* sp. isolates from our collection (Supporting Figure 16). EM imaging showed an ESM composed of a high density network of fibers that were seen to decorate the spore body of *Paenibacillus* sp. spores. The corresponding ‘pearls-on-strings’ 2D class averages confirm that these fibers are indeed F-ENA-2a-like. Although some spores have an exosporal sacculus with a clear crystalline hexagonal pattern (with unit cell dimensions that match the lattice of ExsY sheets^25^), many spores do not have a pronounced exosporium. Interestingly, ENAs seem to directly emerge from the surface of the spore body. Since no exosporium leader sequence motif was detected in the F-Anchor-2a sequence, we hypothesize that the spore coupling mechanism for F-ENA-2a could be different from the ExsF-based coupling seen in *B*. *thuringiensis*.

### F-ENA fibers are also produced by anaerobic bacteria of the class Clostridia

To explore the distribution of the F-ENA fibers in the Bacillota phylum, we performed homology searches for the DUF4183 domain-containing proteins in the non-redundant protein sequence database. The results showed that F-ENA homologs are widely distributed in both the Paenibacillaceae and Bacillaceae families (class Bacilli), and even within the anaerobic bacteria of the class Clostridia. By analyzing F-ENA homologues in various *Clostridia* strains, we show that the mechanism of F-ENA anchoring does indeed seem to vary. For example, in *Clostridium aceticum* strain DSM 1496, the major F-ENA subunit (locus: CACET_c28680; protein: AKL96313.1; hereafter referred to as F-ENA-3) is flanked by CACET_c28690 (protein: AKL96314.1; hereafter referred to as F-Anchor-3) which is composed of one N-terminal DUF4183 domain, four DUF11 domains, one CsxA-like domain, and two additional DUF4183 domains. AF3 predicts a hetero-hexameric complex between 3 copies of F-ENA-3 (Supporting Figure 17a), one copy of F-Anchor-3 and two copies of CsxA (i.e. the major subunit of the *Clostridium* exosporium). As expected, the F-ENA-3 trimer is predicted to dock onto the triad of DUF4183 domains of F-Anchor-3, while the CsxA-like domain of F-Anchor-3 is predicted to form a heterotrimer with the two CsxA copies. Based on this, we hypothesize that the F-ENA-3 pilus is tethered to the *Clostridium aceticum* exosporium via integration of F-Anchor-3 into the CsxA crystal lattice. It is interesting to note that the number of DUF11 domains in F-Anchor-3 tends to vary from zero to four across different homologues within the *Clostridia* class (Supporting Figure 17b).

Consistent with this prediction, we observed F-ENA fibers using cryo-EM on the surface of an anaerobic bacterium of the family Anaerovoracaceae within the class Clostridia (Figure 7, Supporting Figure 18). The fibers, several microns in length, were present on the surface of the spores and exhibited a repeating “beads on a string” pattern characteristic of F-ENA filaments observed in *Bacillus* and *Paenibacillus* spores (Figure 7a, d, e). The averaged power spectrum of aligned raw particles was indexed to estimate all potential helical symmetries (Supporting Figure 18), which were further tested in helical reconstruction. The correct symmetry was determined to be C3, with a rise of 71.2 Å and a twist of 74.4 degrees. The final reconstruction, with helical symmetry applied, reached a resolution of approximately 4.1 Å, as judged by map-to-map FSC. Cα tracing at this resolution was straightforward, and the sequence of the building block was predicted using ModelAngelo to be MDF2656102 (hereafter referred to as F-ENA-4) and the identity of the corresponding gene was confirmed by PCR.

F-ENA-4 exhibits a relatively high sequence identity of 66.9% to F-ENA-3 from *Clostridium aceticum*, but only moderate sequence identities to F-ENA and F-ENA-2a, at 21.5% and 29.1%, respectively. Despite the moderate sequence identities among these three experimentally determined F-ENA fibers, all three share a C3 symmetry and similar packing of the DUF4183 domain trimer (Figure 7b, c). The variation in helical symmetry is primarily due to the differences in the lengths of the N- and C-termini that form spacers between the “beads” composed of the DUF4183 domain trimers (Figures 5f, 7c). In particular, the shortest spacer of 34 Å is present in F-ENA, the largest of 95 Å in F-ENA-2a, and F-ENA-4 falls in the middle with 71 Å. Within this extended region, composed of a six-stranded β-barrel in F-ENA-4, a similar level of hydrogen bonding was observed (Figure 7f) compared to the other two F-ENAs. Furthermore, F-ENA-4 also possesses an inserted N-terminal hook that tightly binds to the next layer of the DUF4183 trimers (Figure 7g). Interestingly, and in contrast to the other two F-ENAs, extra cryo-EM densities, putatively representing post-translational modifications, were observed on eight residues per subunit. These residues, five serines and three threonines, are likely to correspond to *O*-linked glycosylation sites.

In summary, F-ENA fibers appear to be ubiquitous among members of the Bacillota, characterized by a conserved DUF4183 fold and N-terminal hook, yet displaying variability in the surface biophysical properties and inter-domain distances between DUF4183 domains.

## Discussion

Bacteria have a remarkable ability to populate and colonize a myriad of interfaces and surfaces. Under favorable growth conditions, they develop large, actively growing communities that -as resources gradually deplete-transform into structured biofilms, consisting of sessile cells and a complex, multi-component extracellular matrix (ECM) that encases and reinforces the biofilm^26^. Within the biofilm, there is often a small population of persister cells that are either fully metabolically dormant or exhibit selective inactivation of genes, which enables them to survive highly unfavourable, otherwise lethal conditions such as nutrient deprivation, antibiotic stress or exposure to detergents^27^. This specialized survival strategy guarantees persistence of the population when conditions become favourable again. Members of the Firmicutes phylum have, in a sense, perfected the concept of ‘persister cells’ through the development of endospores, a highly specialized dormant cell subtype that is optimized for survival and dissemination. The cell’s ability to produce endospores creates an additional layer of complexity with respect to biofilm formation and allows for fine-tuning of the relative abundances of vegetative cells and endospores in response to environmental stimuli. The impact of the cell-to-spore ratio on biofilm stability, structure, resilience, and adhesive properties is still poorly understood. Reports on biofilms of *B. subtilis* (*Bs*) -the most extensively studied species within the Firmicutes phylum-tend to predominantly focus on cells and the ECM, which for Bs is composed of extracellular DNA, exopolysaccharides, TasA protein fibers and BslA^28^. Nevertheless, it is clear that spores can also be found within *Bs* biofilms^29^. For *Bacillus cereus* (*Bc*), depending on the strain and culture conditions, spores can even constitute up to 90% of the total biofilm population^30^. Biofilms dominated by spores upend the conventional perspective on biofilm formation and call into question the broader applicability of lessons learned from *Bs* biofilms to those formed by *Bc* and *Bt*^31^. Recent cryoEM studies of Bc spores show these are decorated with unique proteinaceous appendages not found on vegetative cells ^14,16^. Here, we expand the family of endospore appendages and find they can give rise to a dense extrasporal matrix distinct in composition to the extracellular matrix surrounding biofilms dominated by vegetative cells.

In the conditions tested in the current work, biofilms of *B. thuringiensis* serovar Kurstaki were predominantly (94%) composed of spores and toxin crystals, with only a small minority of cells being present. We observed the presence of an ESM that was dominated by spore attached endospore appendages. Using cryoEM, we resolved the structure of the S-type variant, which was found to be structurally closely related to the S-type ENAs observed on *B. paranthracis* (*Bp*) NVH 0075-95. Moreover, we identified a previously unknown ENA sub-type: flexible, 5 nm diameter ENAs (F-ENA) coupled to the exosporium of *Btk*. The unique identity of the F-ENA protomers was determined through cryo-identification. F-ENAs are composed of small (10 kDa) DUF4183 proteins that trimerize and stack axially into a helical ultrastructure. We showed that F-ENAs are (i) tethered to the exosporium via a collagen-like protein, F-Anchor, which docks onto ExsF via an N-terminal exosporium leader sequence, and (ii) serves as an assembly platform for F-ENA formation by displaying a C-terminal DUF4183 domain on a long collagen triple helix that pierces through the hairy nap layer (Figure 6). At the spore distal terminus, F-ENAs are decorated with a flexible tip fibrillum that consists of the collagen-like protein F-BclA. Although F-BclA has a variable number of collagen triplet repeats, its C-terminal C1q-like domain is highly conserved throughout the *B*. *cereus* group.

**Figure 6.**
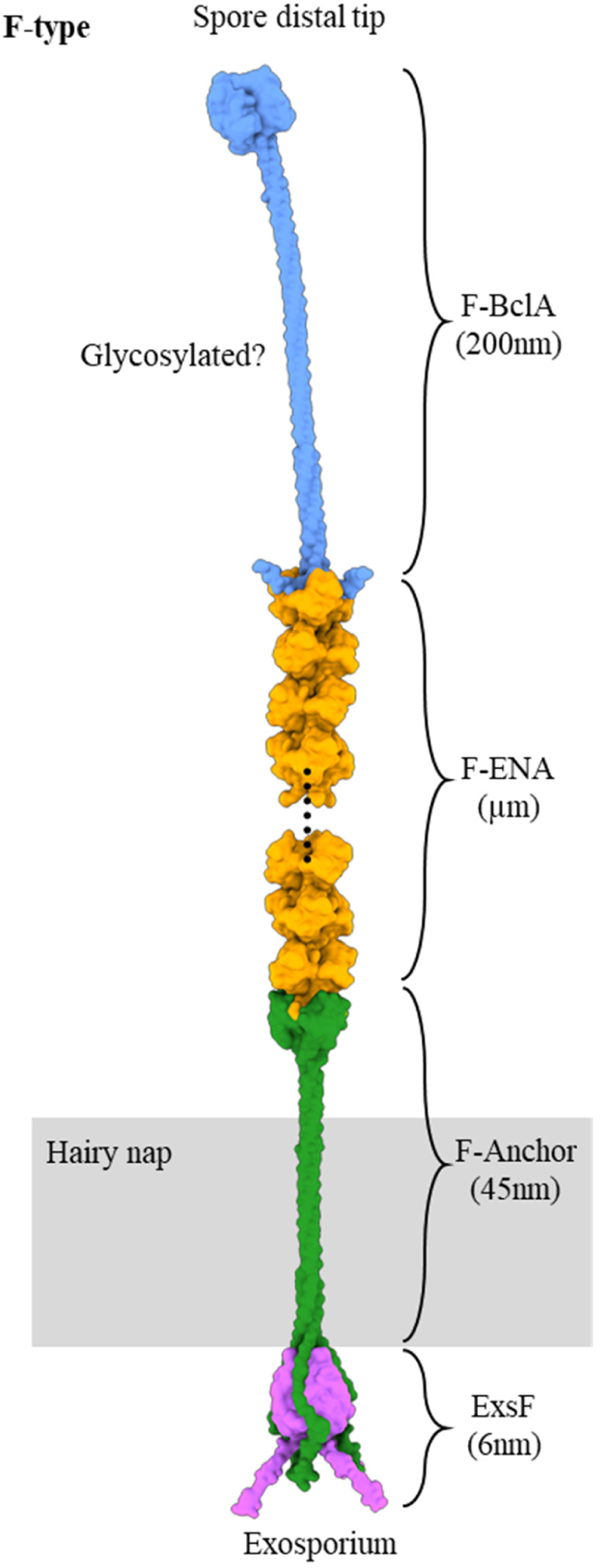
Composite molecular model of F-type endospore appendages found across the *Bacillus cereus* sensu lato group. At the spore proximal terminus F-ENA is docked onto the exosporium basal layer via the F-ENA / F-Anchor / ExsF ternary complex. At the spore distal terminus, the F-ENA tip fibrillum is composed of the collagen-like protein F-BclA.

**Figure 7.**
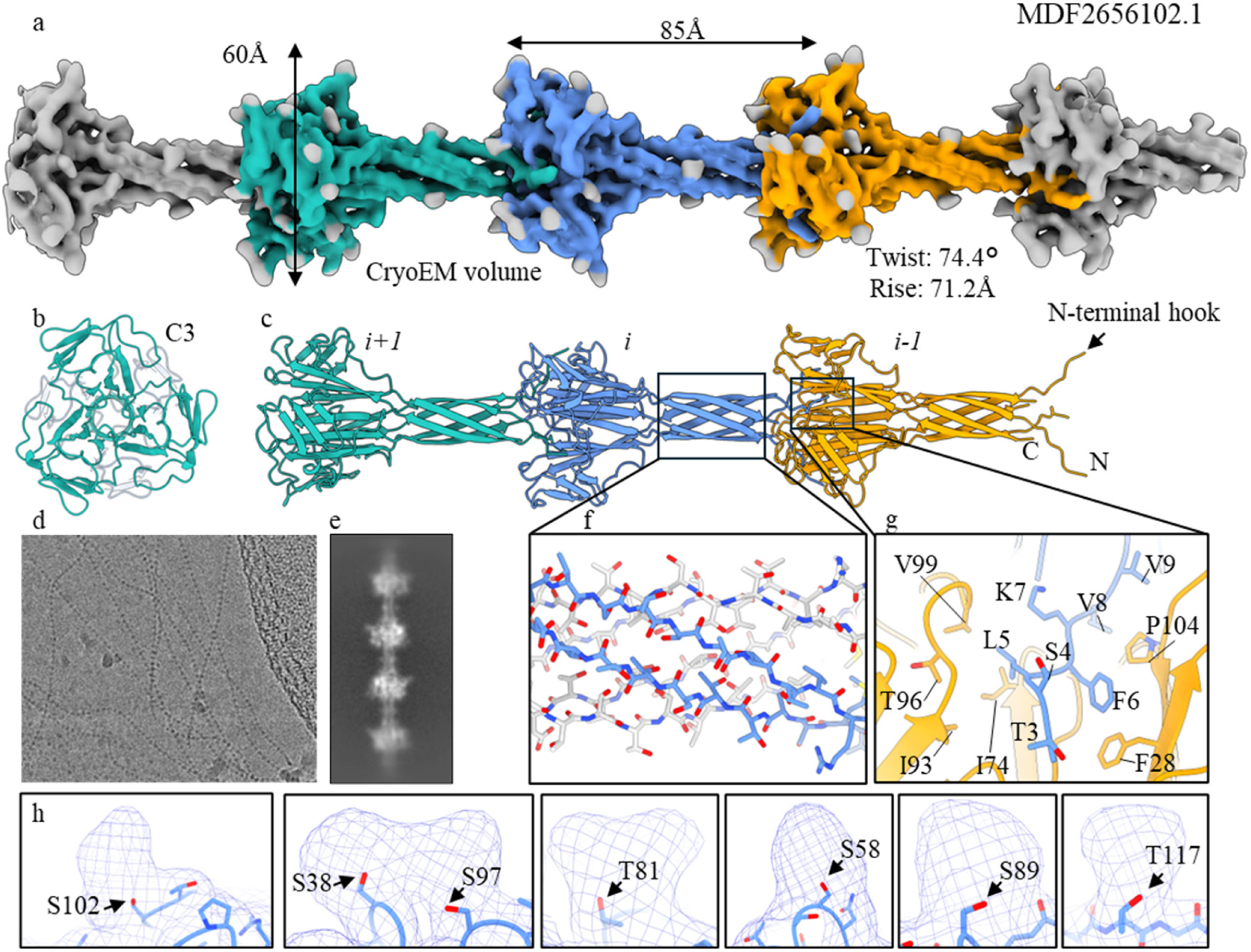
CryoEM of F-type-3 ENA fibers of an uncultivated member of the family Anaerovoracaceae within the class Clostridia. (a) reconstructed cryoEM volume with (b) and (c) the corresponding top and side view of the molecular model shown in cartoon representation; (d) cryoEM micrograph of F-ENA-3 fibers with corresponding 2D class average in (e); (f) zoom-in of the beta-cylinder extension; (g) docking site of the N-terminal hook; (h) Unmodelled densities corresponding to putative O-glycosylation sites.

Our findings suggest that F- and S-ENA likely both contribute to the clustering of *Btk* spores. In a previous contribution, we showed that spore aggregation in *Bp* was dependent on the L-ENA tip fibrillum protein L-BclA, which is localized on the termini of both S- and L-ENA^23^. Given that *Btk* does not produce L-ENA fibers and lacks a genomic copy of *l-bcla*, it is clear that *Btk* aggregation is not driven by L-BclA. We hypothesize that F-BclA could have adopted a role similar to L-BclA, in that it might bind to (a specific receptor on) the surface of the spore to mediate aggregation. The coupling mechanism of F-BclA onto F-ENA mimics the homotypic axial interactions that are unique to the F-ENA pilus. Consequently, F-BclA binding is expected to be limited to F-ENA pili, i.e. with no promiscuous binding to different ENA subtypes. Although the exact identity of the S-ENA ruffle protein(s) remains unclear, a HMMER search using L-BclA as a query against the *Btk* genome yields AGE80398.1. This is a collagen-like protein with (i) a C-terminal C1q domain that has 47.89% sequence identity to the C1q domain of L-BclA, and (ii) with a cysteine-rich (5 cys) N-terminus that carries a signature CC-motif. A CC motif is also found in the S- and L-ENA protomers, as well as the N-terminal connector of L-BclA. Notably, this protein does not contain an exosporium-leader sequence. Whether or not AGE80398.1 constitutes the tip-fibrillum of S-ENA, will be the subject of further study.

Regardless of the exact molecular mechanism of binding, one could ask why *Btk* spores have developed independent, seemingly redundant methods to self-aggregate. The widespread distribution of S and F-ENA operons across the *Bacillus cereus* group and the strong conservation of F-BclA suggest spore aggregation could be a conserved trait of *Bc* group spores. Functional redundancy in the form of the production of various ENA subtypes, each decorated with distinct tip-fibrillae, could be a logical adaptation to the evolutionary pressure to maintain the ability to aggregate. Moreover, S-ENA and F-ENA may adopt slightly different roles in cellular clustering. S-ENA fibers are long (multiple micrometers) and relatively rigid compared to F-ENAs, making them more likely to maintain long-range spore contacts. Their structure enables them to grapple spores that lie beyond the first sphere of nearest neighbours within a spore cluster. In contrast, F-ENAs are more flexible and remain distributed closer to the spore surface. As such, we anticipate that F-ENA will predominantly couple directly neighbouring spores. A similar division of labor between establishing long-range versus short-range contacts is at play for *Bp,* where L-ENA fibers typically measure only a few 100 nm or less. How the differences in molecular architecture, tensile strength and flexibility of F-, L- and S-ENA translate to the macroscopic material properties of the biofilm remains to be studied.

It is remarkable that *Bs* spores are devoid of ENAs. Spore clustering could provide a fitness advantage in numerous defensive scenarios, such as protection against predatory pressures (e.g. amoeba grazing), UV radiation (unlikely for the rhizosphere habitat of *Bs*, but potentially relevant in dispersion), macrophage phagocytosis, etc. Having said that, in a cell-rich biofilm setting, spore clustering may not be a requirement, because the ECM, in particular TasA fibers secreted by the cells^32^, will also encase the spores embedded within the biofilm, thereby abrogating the necessity for direct inter-spore coupling. For biofilms that are dominated by spores with minimal or no ECM present, however, ENAs likely take up a compensatory functional role in lieu of the ECM. Secondly, many of the species in the *Bacillus cereus* group sensu lato exhibit pathogenic lifestyles. ENA-mediated spore clustering could factor into offensive strategies by facilitating the formation of spore aggregate sizes that approach or exceed the minimal infectious dose for a given host. If indeed proven to be the case, S- and F-ENA could become valuable targets to increase the virulence of Bt and with it the efficacy of Bt-based biopesticides.

Including the newly identified F-ENA, the number of characterized ENA-subtypes stands at three – a number that is expected to increase as research continues. S-, L- and F-ENAs share little sequence identity, are all architecturally distinct and are tethered to different spore surfaces: S-ENAs are associated with the spore coat^14^, while L- and F-ENAs are anchored to the exosporium^16^). The common denominator across this group of ENAs is that they all terminate in a collagen-like tip fibrillum. This suggests that the main biological function of ENAs is to serve as a structural platform to display C1q-like domains at their distal ends, which have presumed adhesive properties. The F-ENA-2a type ENAs found in the Paenibacillacea described in this contribution deviate from that pattern in that we did not observe any tip-fibrillae via nsTEM, nor did we identify putative ruffle candidates through genetic screening. However, we did observe antiparallel, lateral co-alignment of F-ENA-2a fibers in nsTEM. Importantly, we did not observe such bundling in cryoEM, suggesting that F-ENA-2a bundling predominantly occurs under dehydrated conditions. In natural settings, spores can be exposed to desiccation events, and hence ENA bundling might be a response to dehydration. (Anti-parallel) fiber bundling due to lateral interactions, as seen with F-ENA-2a, has been observed for TasA in *B. subtilis*^33^, archaeal bundling pili (ABP) in *Pyrobaculum calidifontis*^34^ and more recently in ABP-like fibers in *Pyrodictium abyssi* (ABPx)^35^. In these cases, the bundling likely contributes to the stability of their respective ECMs. TasA, ABP and ABPx are all donor-strand exchanged fibers. Although F-ENA-2a falls outside that structural category, it may have evolved to achieve a similar goal, i.e. to bridge contacts between spores, either through entropic entanglement or direct fiber-fiber coupling.

---

**Supporting Figure 1.**
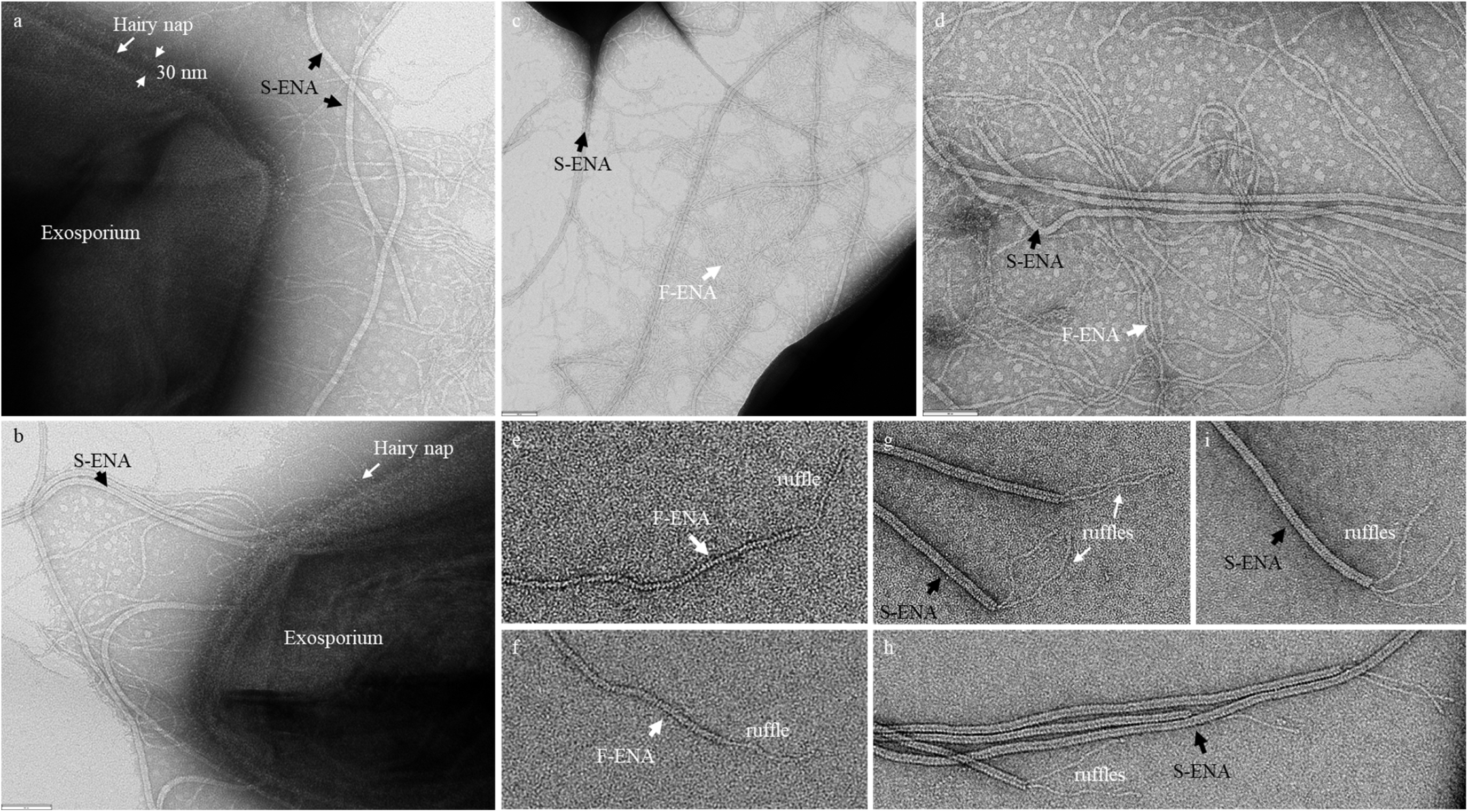
nsTEM micrographs of *Bacillus thuringiensis* subsp. kurstaki endospore appendages. (a) Measurement of the thickness of the hairy nap layer (±30nm). (b) S-ENA and F-ENA fibers are seen to decorate the spore surface, with F-ENA coupled specifically to the exosporium. (c) and (d) The region between two neighbouring spores. (e) and (f) Singular tip fibrilla that decorate the spore distal terminus of F-ENA. (g) – (i) One to three tip fibrilla that decorate the spore distal terminus of S-ENA.

**Supporting Figure 2.**
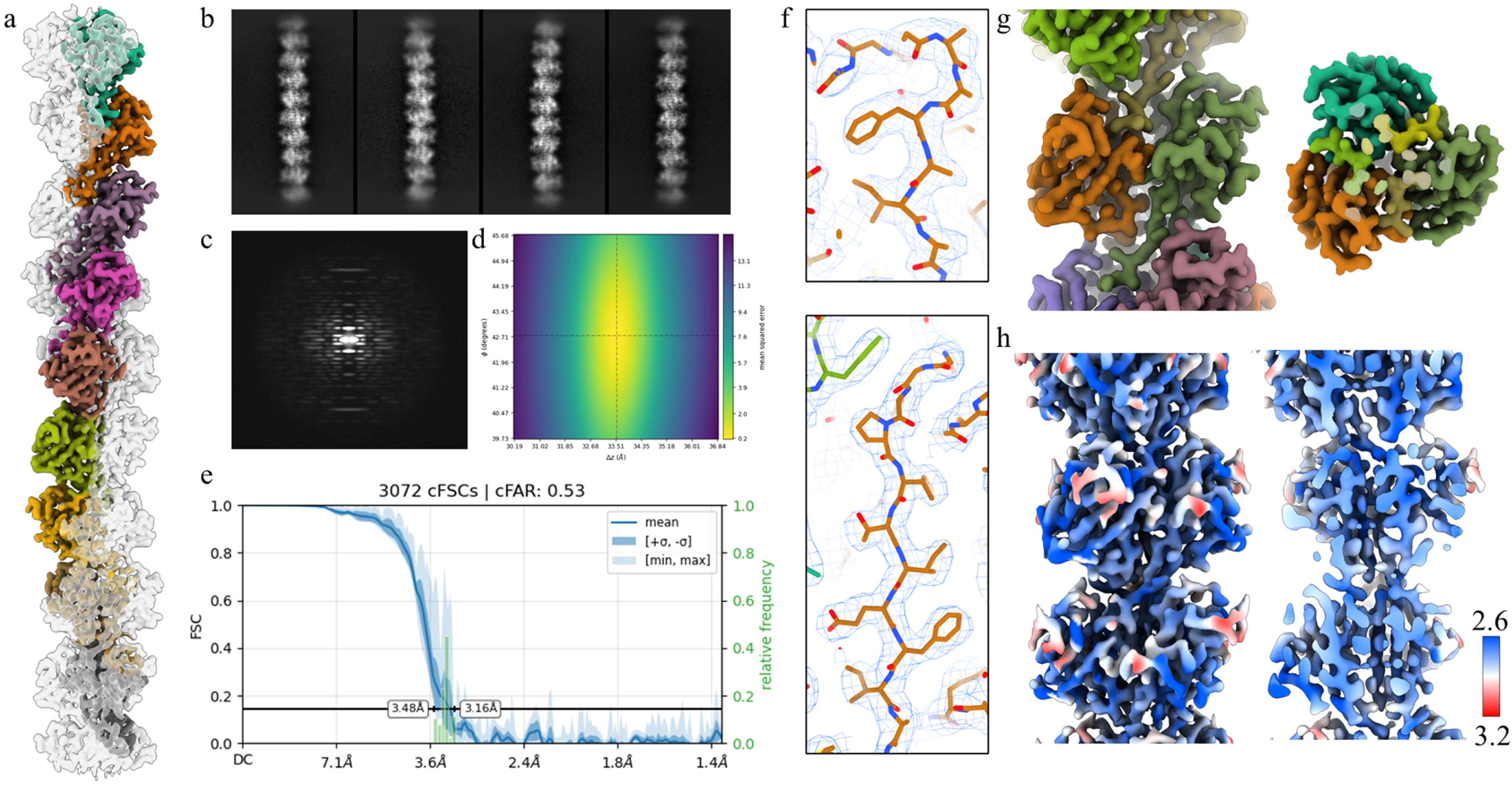
Helical reconstruction of *ex vivo* F-ENA pili. (a) Reconstructed cryoEM volume with one protofibril colour-coded according to chain ID; (b) 2D class averages: vertical box size measures 358Å; (c) Power spectrum; (d) Helical symmetry error surface with a minimum at 42.73 degrees and 33.54Å; (e) conical Fourier Shell Correlation curve produced by Orientation Diagnostics job in CryoSPARC (v4.6.2); (f) Illustrated segments are used to represent the map quality; (g) Zoom-in of a single fiber segment in a side (left) and on-axis (right) view (EMReady map); (h) Surface colouring according to local resolution. Map generated by Local Filtering in CryoSPARC.

**Supporting Figure 3.**
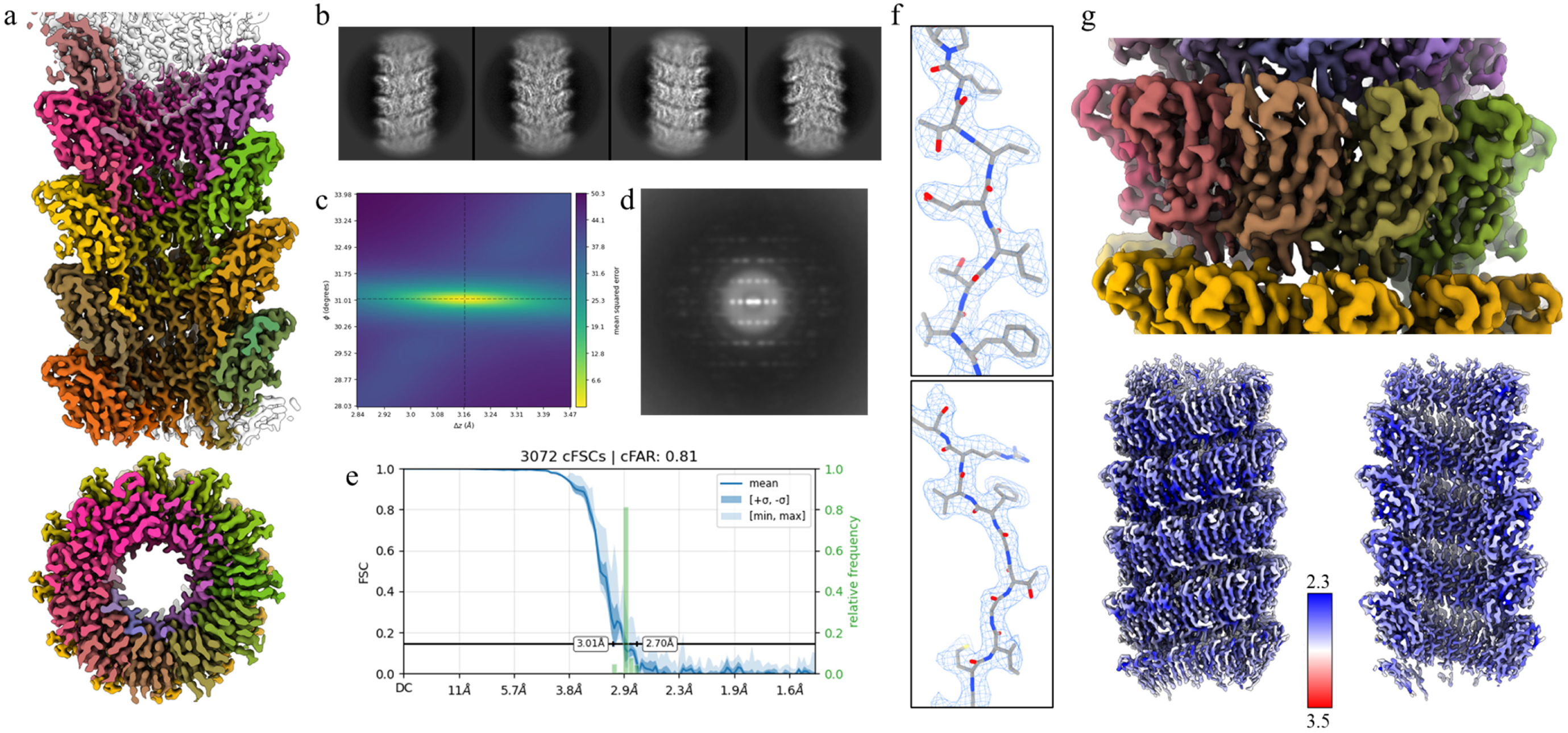
Helical reconstruction of *rec*-S-ENA pili. (a) Reconstructed cryoEM volume of recombinant Ena2A; (b) 2D class averages: box size measures 210Å; (c) Power spectrum; (d) Helical symmetry error surface with a minimum at 31.03 degrees and 3.16Å; (e) conical Fourier Shell Correlation curve produced by Orientation Diagnostics job in CryoSPARC (v4.6.2); (f) Illustrated segments are used to represent the map quality; (g) Zoom-in onto a single helical turn; (h) Surface colouring according to local resolution. Map generated by Local Filtering in CryoSPARC.

**Supporting Figure 4.**
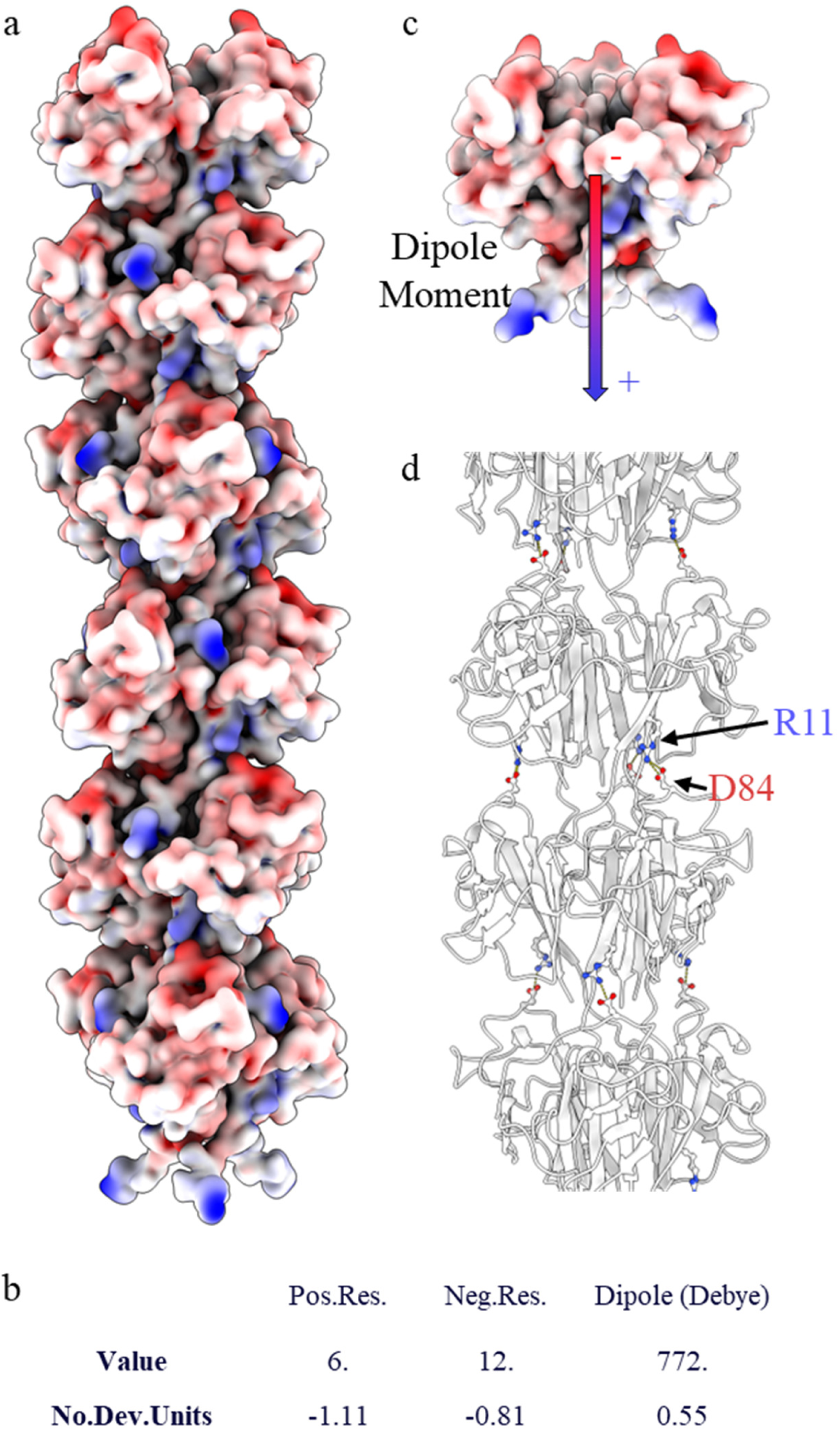
Surface electrostatics and dipole moment of F-ENA. (a) F-ENA pilus surface colour-coded according to electrostatic potential using ChimeraX; (b) Dipole moment of an F-ENA trimer as calculated by the Protein Dipole Moments Server (https://dipole.proteopedia.org/) and visualized onto a single F-ENA trimer (c); (d) Salt bridges between R11 and D84 that are reinforcing the interaction between two successive F-ENA fiber segments are shown in stick representation.

**Supporting Figure 5.**
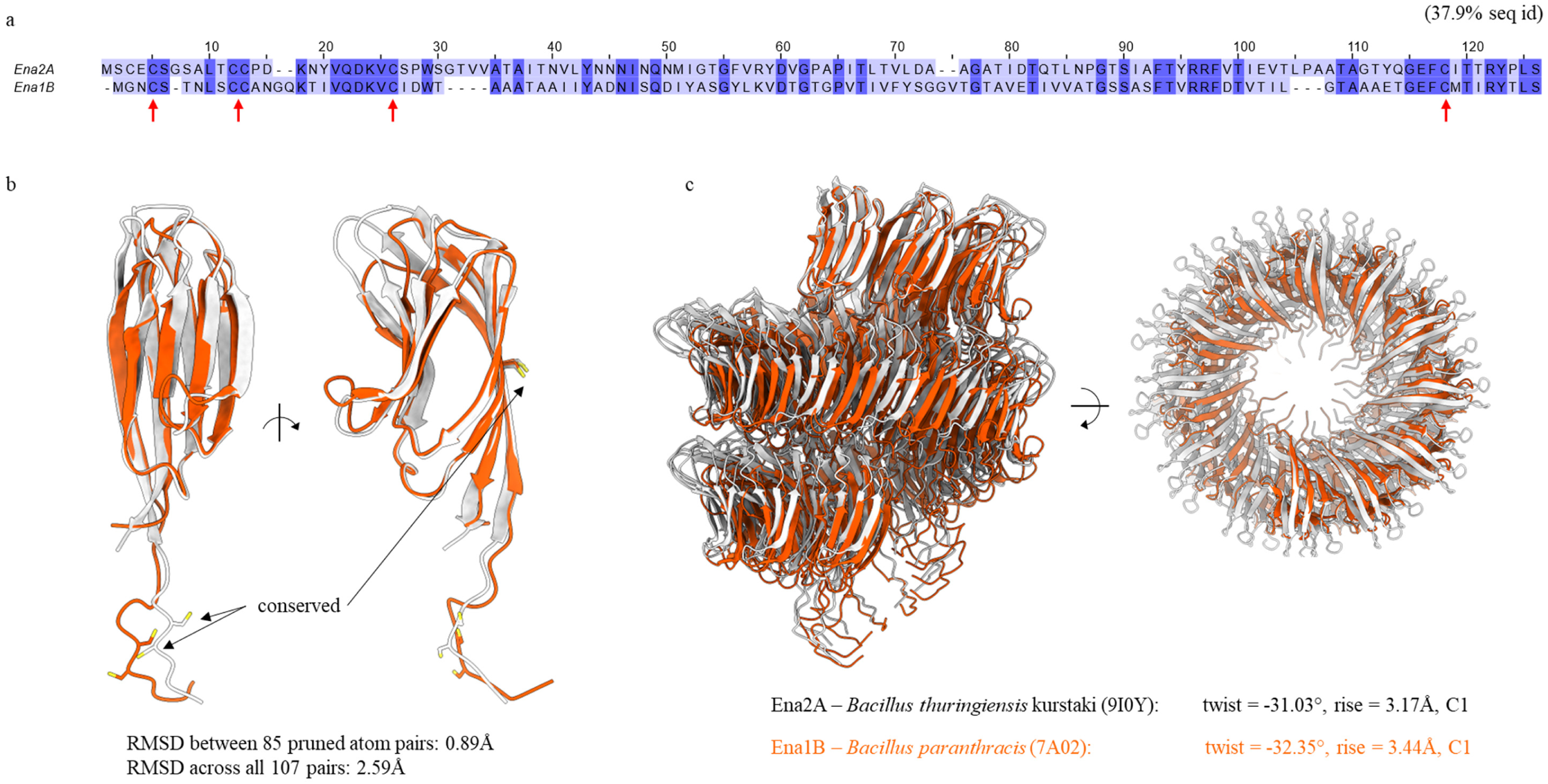
Structural comparison between Ena2A and Ena1B. (a) Pairwise sequence alignment of Ena2A and Ena1B. The sequence identity is 37.9%. Red arrows indicate cysteine positions involved in inter-molecular disulfide bridge formation (b) Structural alignment of the Ena2A (this study) and Ena1B (7A02) monomer (pruned / all atom pair RMSD is 0.89Å / 2.59Å) shows strong conservation of the fold, as well as conservation of the cysteine positions. (c) Structural alignment of the Ena2A and Ena1B fibers highlighting the conserved overall helical ultrastructure.

**Supporting Figure 6.**
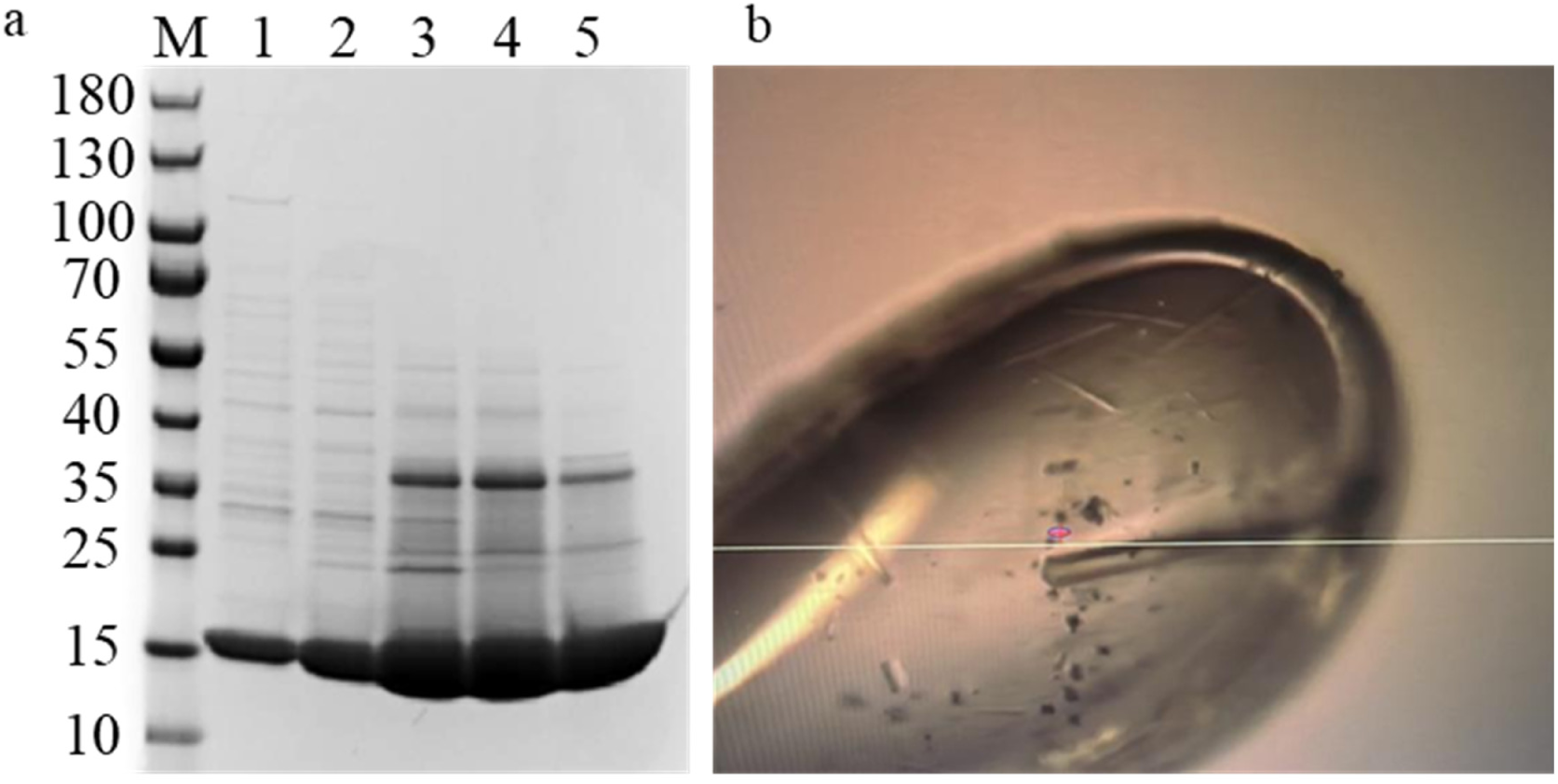
Purification and crystallization of the ExsF/FA_14-35_ complex. (a) SDS-PAGE analysis of ExsF fractions collected during size-exclusion chromatography using a HiPrep 26/60 Sephacryl S-200 HR column. The theoretical molecular weight of the ExsF monomer and trimer is 16.2 and 48.5 kDa, respectively; (c) Vitrified ExsF/FA_14-35_ crystal suspended in a crystallization loop. Crystals were formed in 5 % v/v T-mate pH 7, 0.1 M sodium cacodylate pH 5.3, 15 % v/v PEG smear broad and 10 % v/v ethylene glycol, in the BCS screen (Molecular Dimensions).

**Supporting Figure 7.**
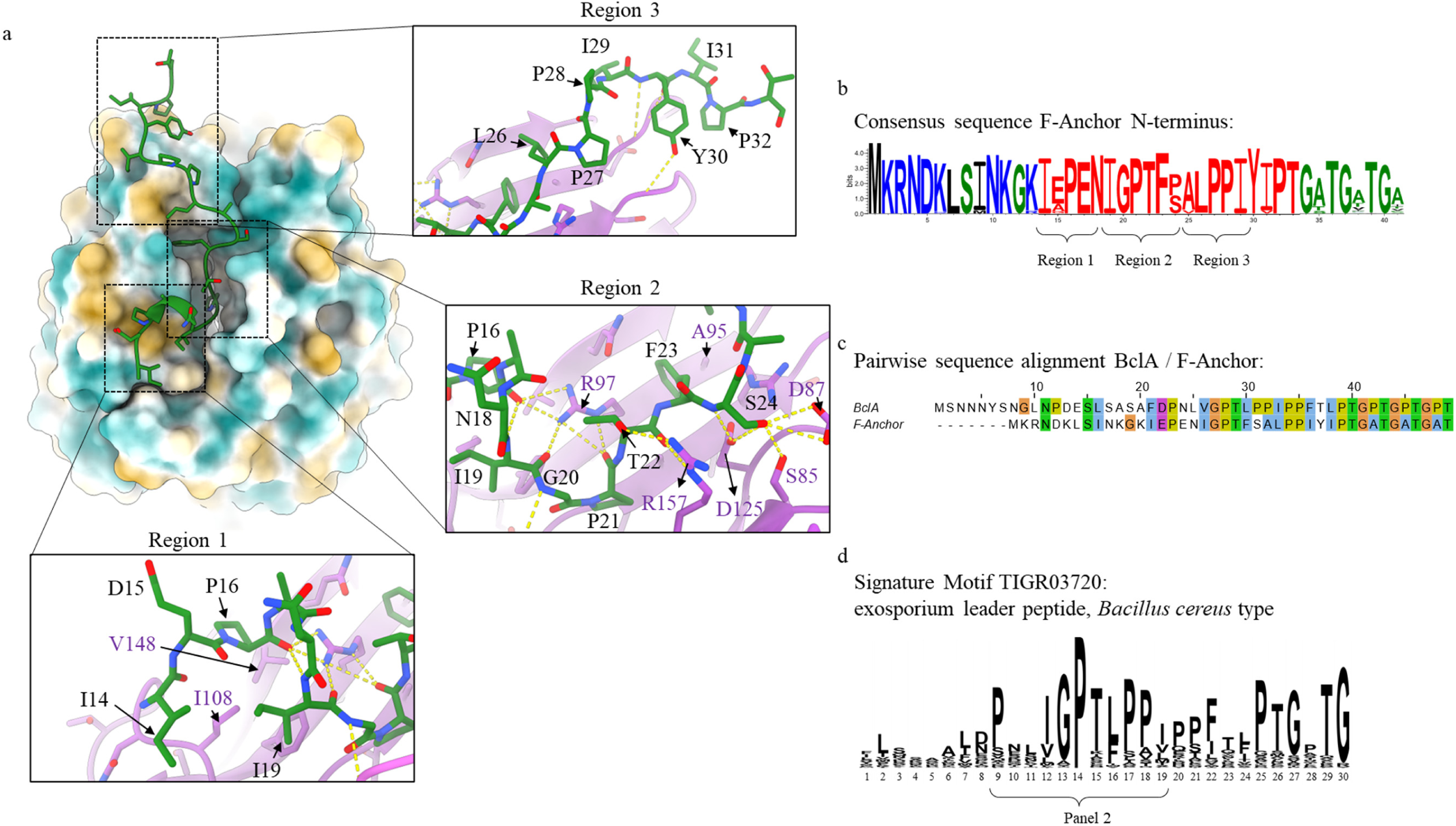
Crystal structure of the ExsF trimer in complex with the exosporium leader peptide of F-Anchor. (a) Interaction interface between the ExsF dimer epitope and F-Anchor_14-28_, identifying three interaction regions. (b) Consensus sequence of the F-Anchor N-terminal region highlights conservation of the exosporium leader sequence. (c) Pairwise sequence alignment between the N-termini of BclA and F-Anchor. (d) Sequence logo of the exosporium leader peptide determined from TIGRFAM entry TIGR03720.

**Supporting Figure 8.**
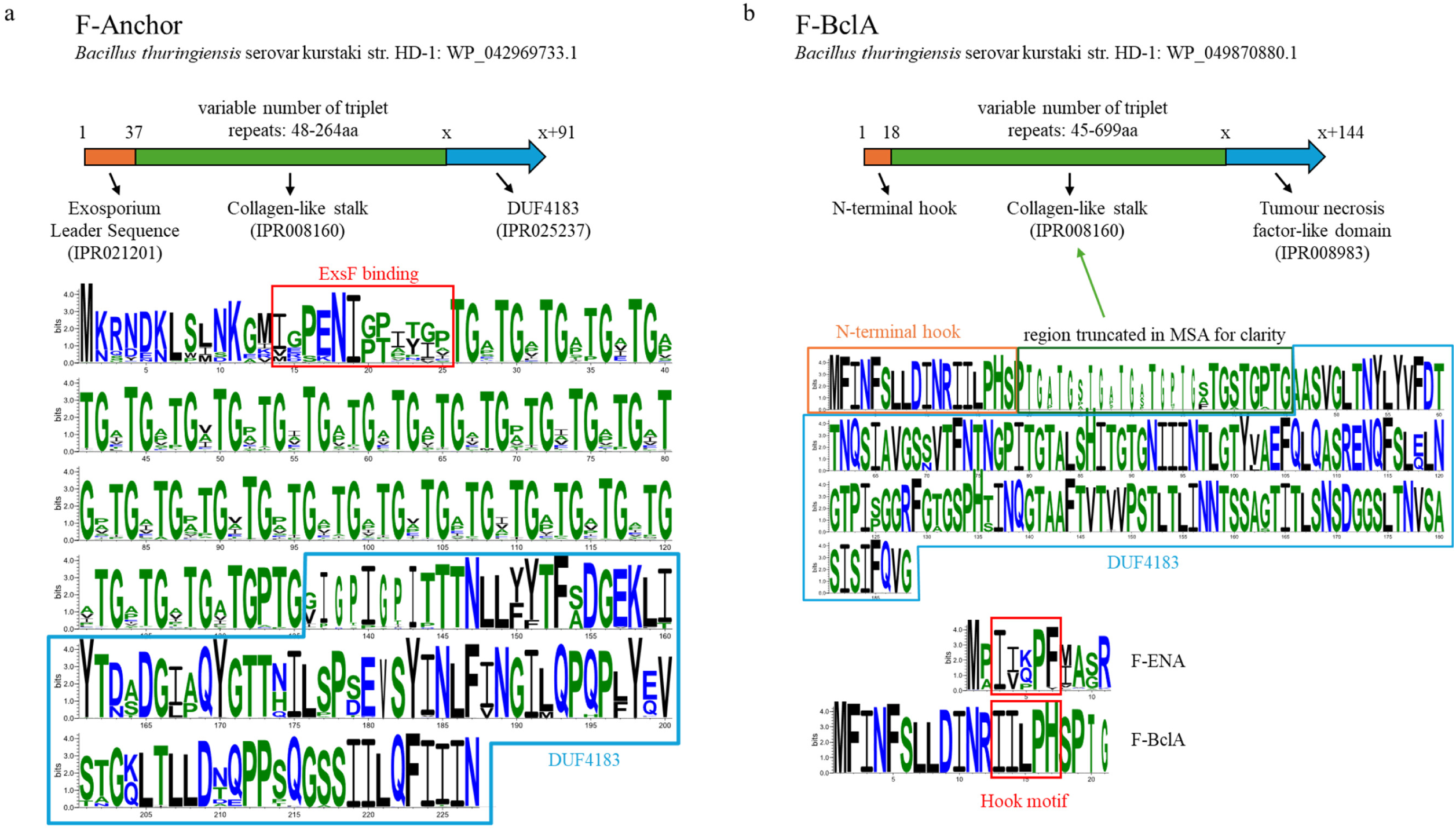
Gene domain organization and sequence conservation of F-Anchor and F-BclA. (a) Primary sequence architecture of F-Anchor composed of an N-terminal exosporium leader sequence (IPR021201) for docking onto ExsF, a collagen triplet-repeats region and a C-terminal DUF4183 domain for coupling onto F-ENA. (b) Primary sequence architecture of F-BclA composed of an N-terminal hook motif for docking onto F-ENA, a collagen triplet-repeats region and a C-terminal BclAC-like domain (IPR008983). (c) Comparison of the F-ENA and F-BclA hook motifs.

**Supporting Figure 9.**
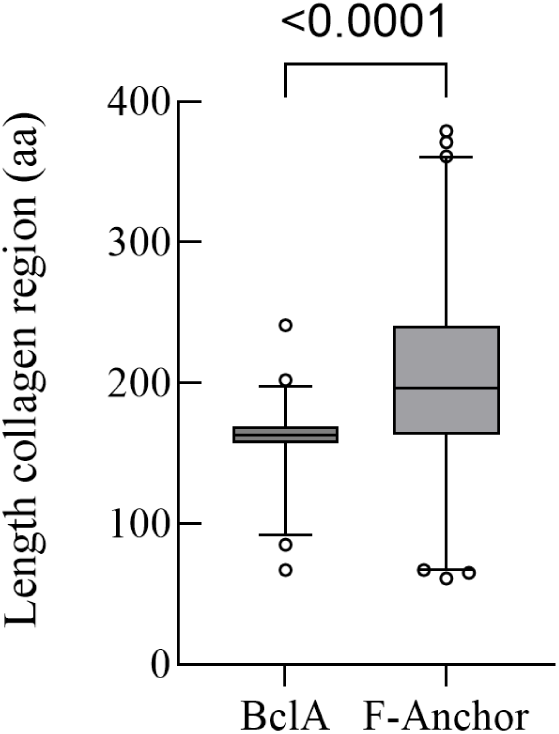
Length distribution of the collagen-like stalk of BclA and F-Anchor across the *Bacillus cereus* sensu lato group. Mean for BclA: 162nm (n=233, box: median, 25^th^ and 75^th^ percentile, whiskers: 1 and 99 percentile), mean for F-Anchor: 201nm (n=301, box: median, 25^th^ and 75^th^ percentile, whiskers: 1 and 99 percentile).

**Supporting Figure 10.**
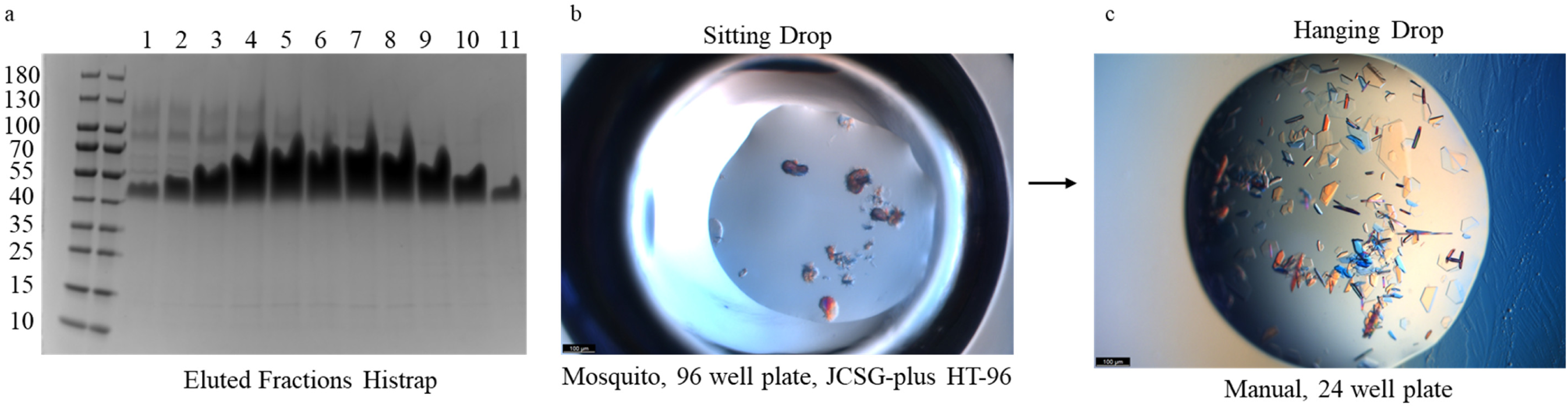
Purification and crystallization of F-BCLA-CTD. (a) SDS-PAGE analysis of F-BCLA-CTD fractions collected during elution of a 5 mL Histrap Fast Flow (Cytiva) column.The theoretical molecular weight of the F-BCLA-CTD monomer and trimer is 16.6 and 49.8 kDa, respectively; (b) Initial crystallization hit obtained in the JCSG-plus HT-96 screen condition C2 in sitting drop geometry, set up using the Mosquito crystallization robot; (c) Crystallization conditions were optimized manually in a hanging drop setup. Plate-like crystals appeared within one week in 1M LiCl, 0.1M citrate pH 4.0, 20% (w/v) PEG 6000).

**Supporting Figure 11.**
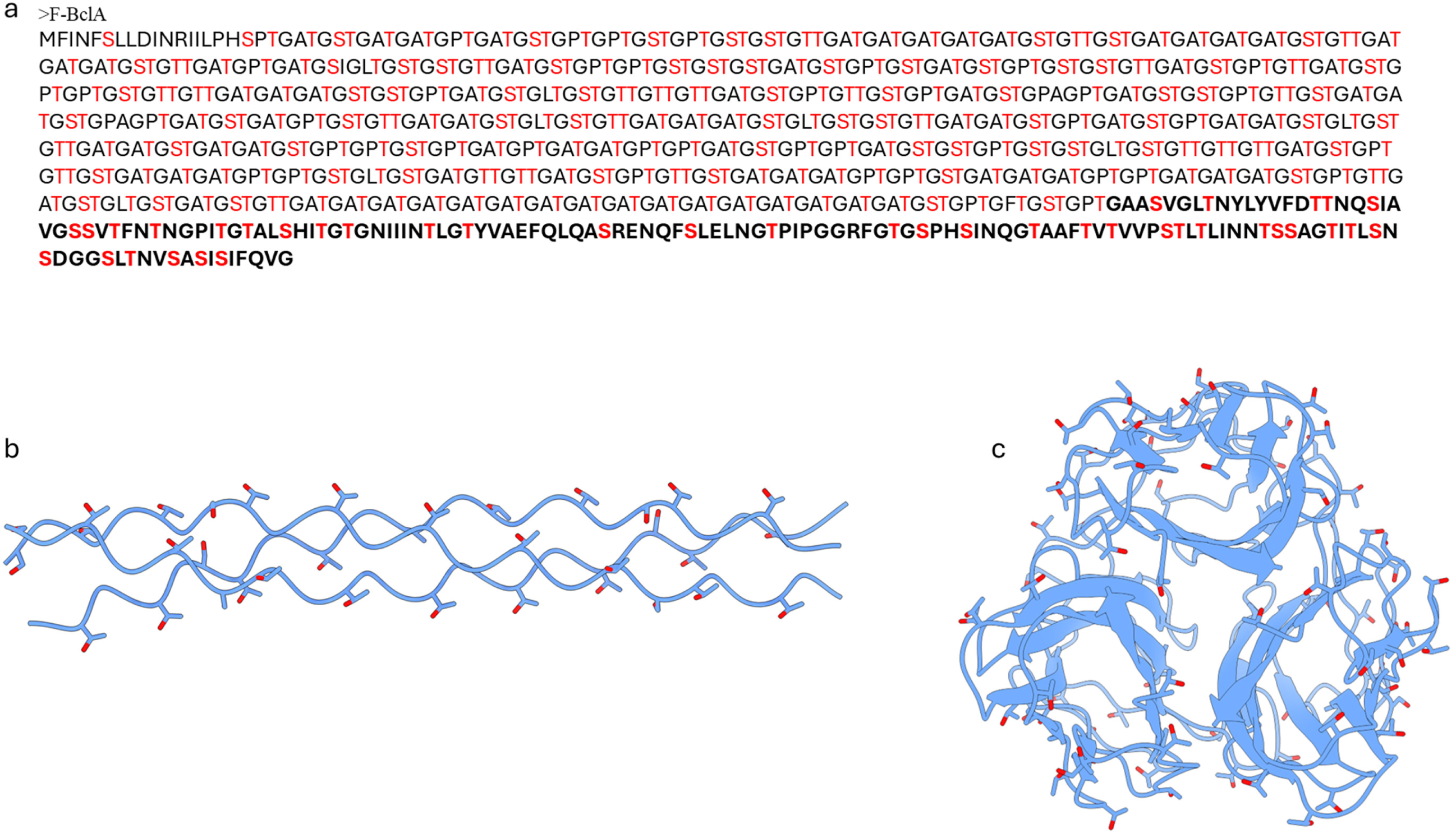
Surface exposed Ser and Thr residues in F-BClA. (a) Primary sequence of F-BclA with S and T residues highlighted in red. The CTD domain is shown in bold; (b) Surface exposed Ser and Thr residues on a segment of the AF3 model of the collagen-like stalk; (c) Surface exposed Ser and Thr residues on the X-ray structure of F-BclA-CTD.

**Supporting Figure 12.**
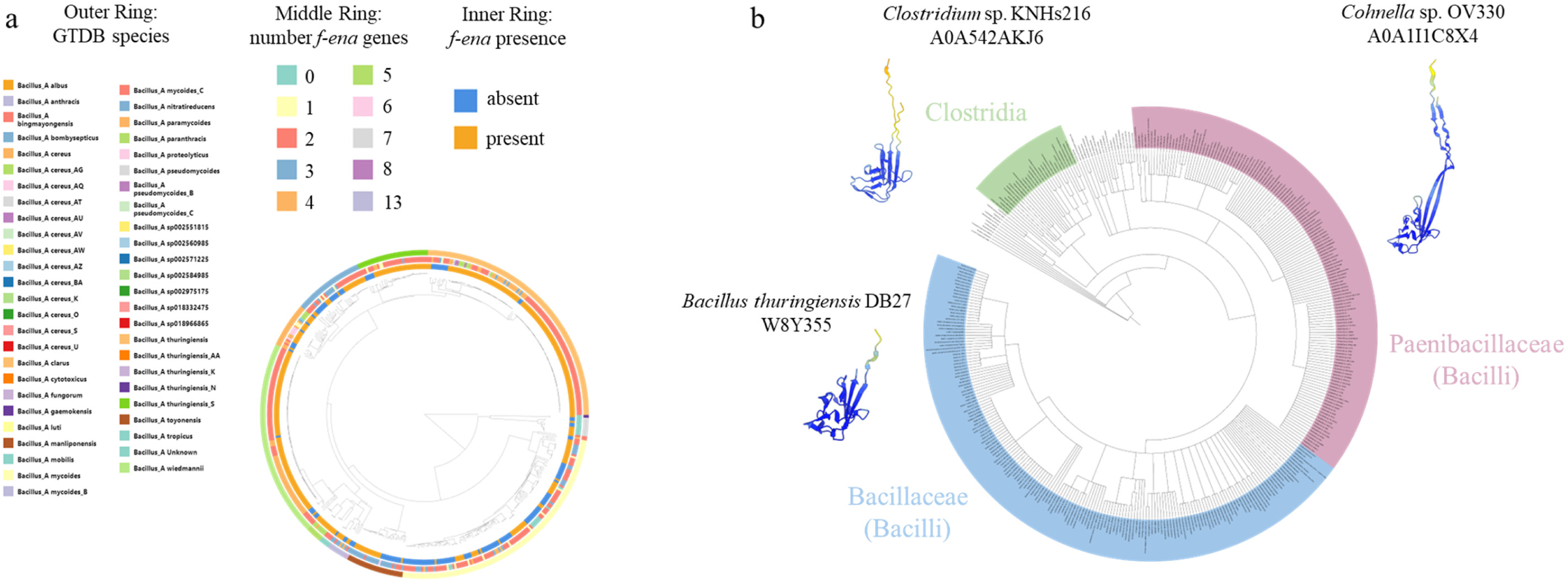
Phylogenetic distribution of F-ENA-like genes. (a) across the *Bacillus cereus* sensu lato group (Btyper database). The inner ring shows the presence or absence of *f-ena* homologues genes, the middle ring the number of *f-ena*-like genes and the outer ring the GTDB species. An interactive version is accessible at https://microreact.org/project/8vm5uYdv6K9XnmnmuuSzE8-btyperdb-f-ena; (b) in the Bacillota phylum (taxid: 1239). AF2 predictions of selected members for each phylogenetic clade only serve illustrative purposes and are not fully representative of the structural diversity within each clade.

**Supporting Figure 13.**
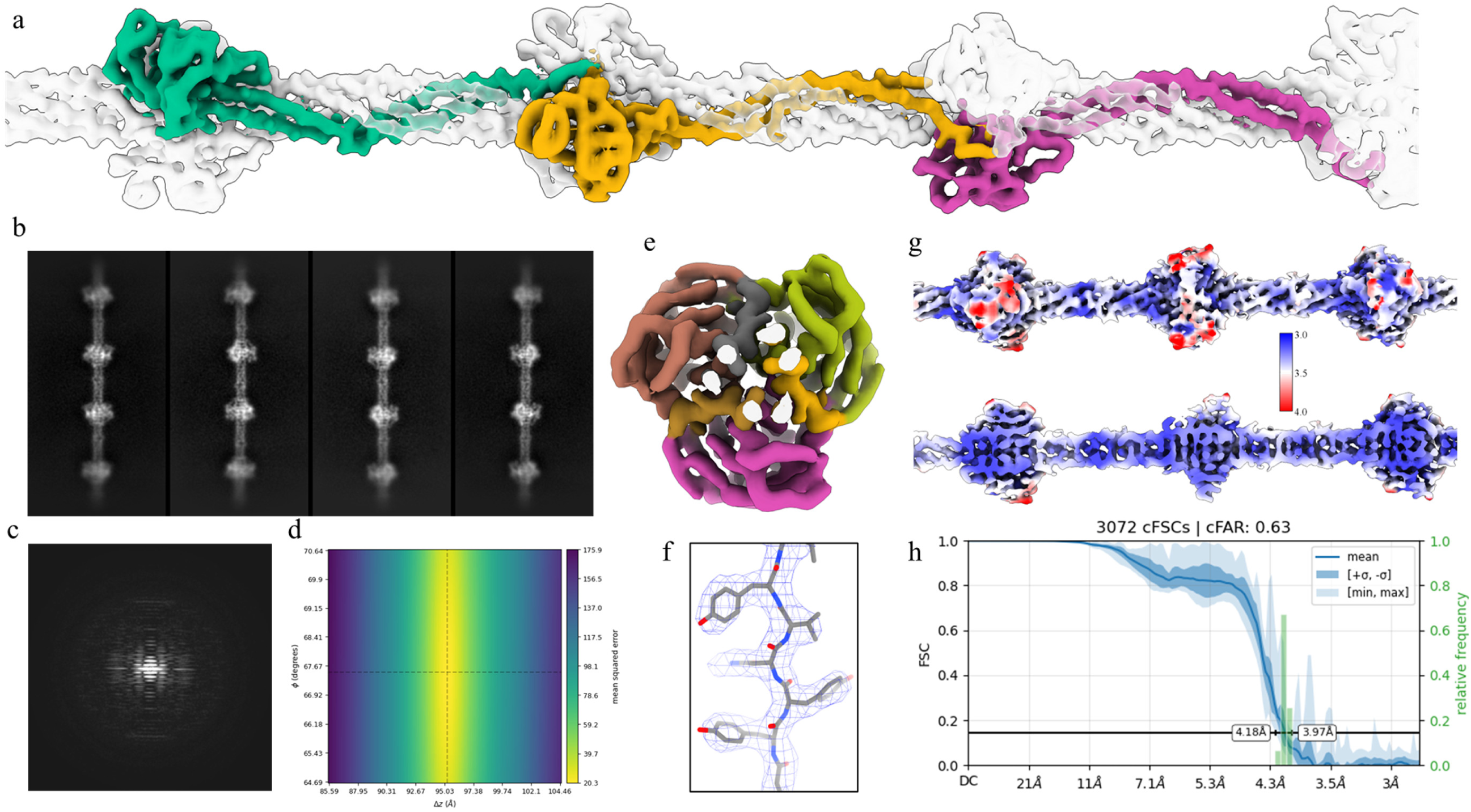
Helical reconstruction of *rec*-F-ENA-2a pili. (a) Reconstructed cryoEM volume of recombinant F-ENA-2a fibers (EMready sharpened); (b) 2D class averages: box size measures 420Å vertically; (c) Power spectrum; (d) Helical symmetry error surface with a minimum at 67.50 degrees and 95.25Å; (e) Top view cross section; (f) F-ENA-2a segment docked in EMReady sharpened map; (g) Surface colouring according to local resolution. The map was generated by Local Filtering in CryoSPARC; (h) conical Fourier Shell Correlation curve produced by Orientation Diagnostics in CryoSPARC (v4.6.2).

**Supporting Figure 14.**
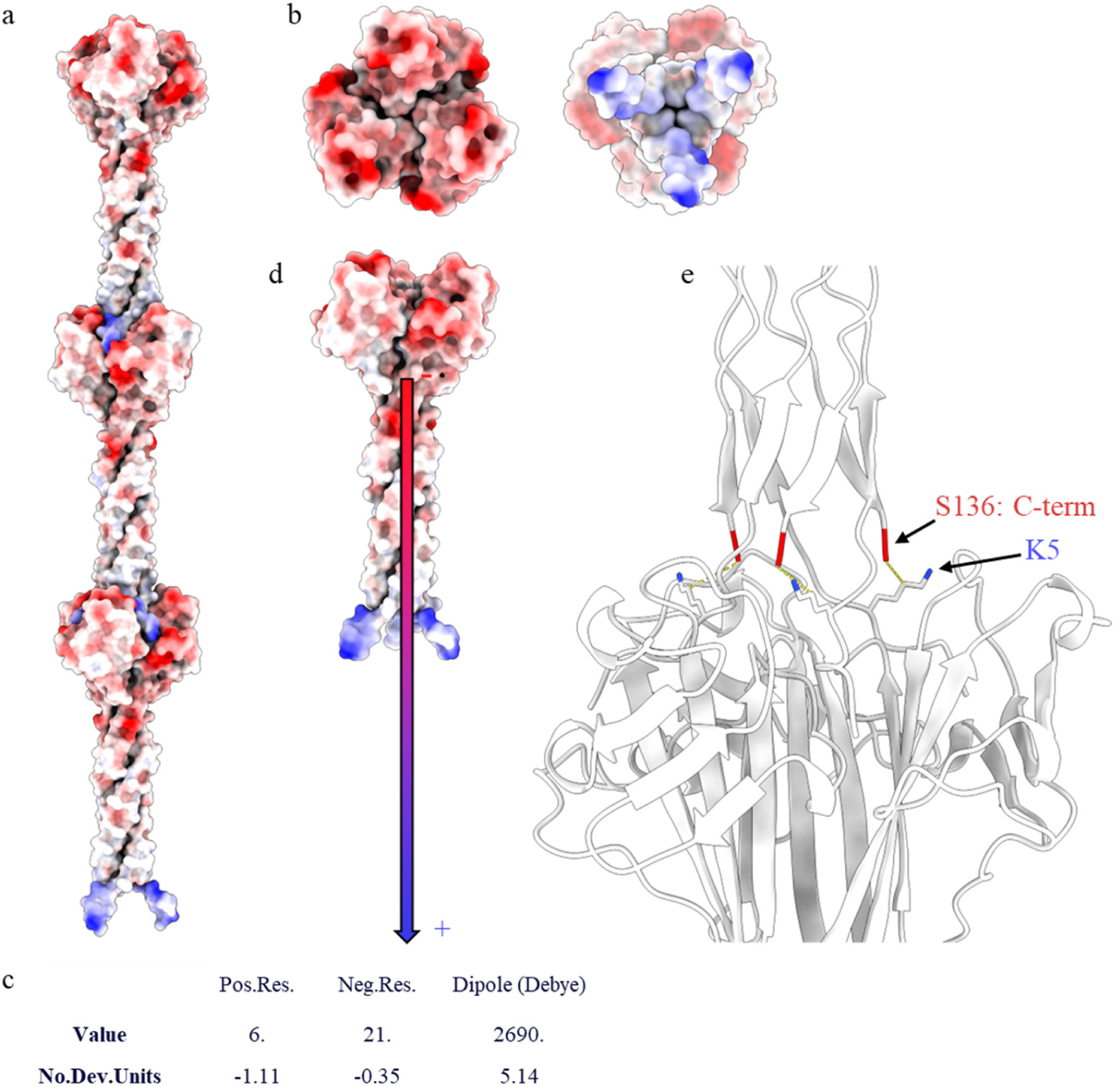
Surface electrostatics and dipole moment of F-ENA-2a. (a) F-ENA-2a pilus surface colour-coded according to electrostatic potential using ChimeraX; (b) Top and Bottom view; (c) and (d) Dipole moment of an F-ENA-2a trimer as calculated by the Protein Dipole Moments Server (https://dipole.proteopedia.org/) and visualized onto a single F-ENA-2a trimer; (e) Salt bridges between K5 and the C-terminus that are reinforcing the interaction between two successive F-ENA-2a fiber segments are shown in stick representation.

**Supporting Figure 15.**
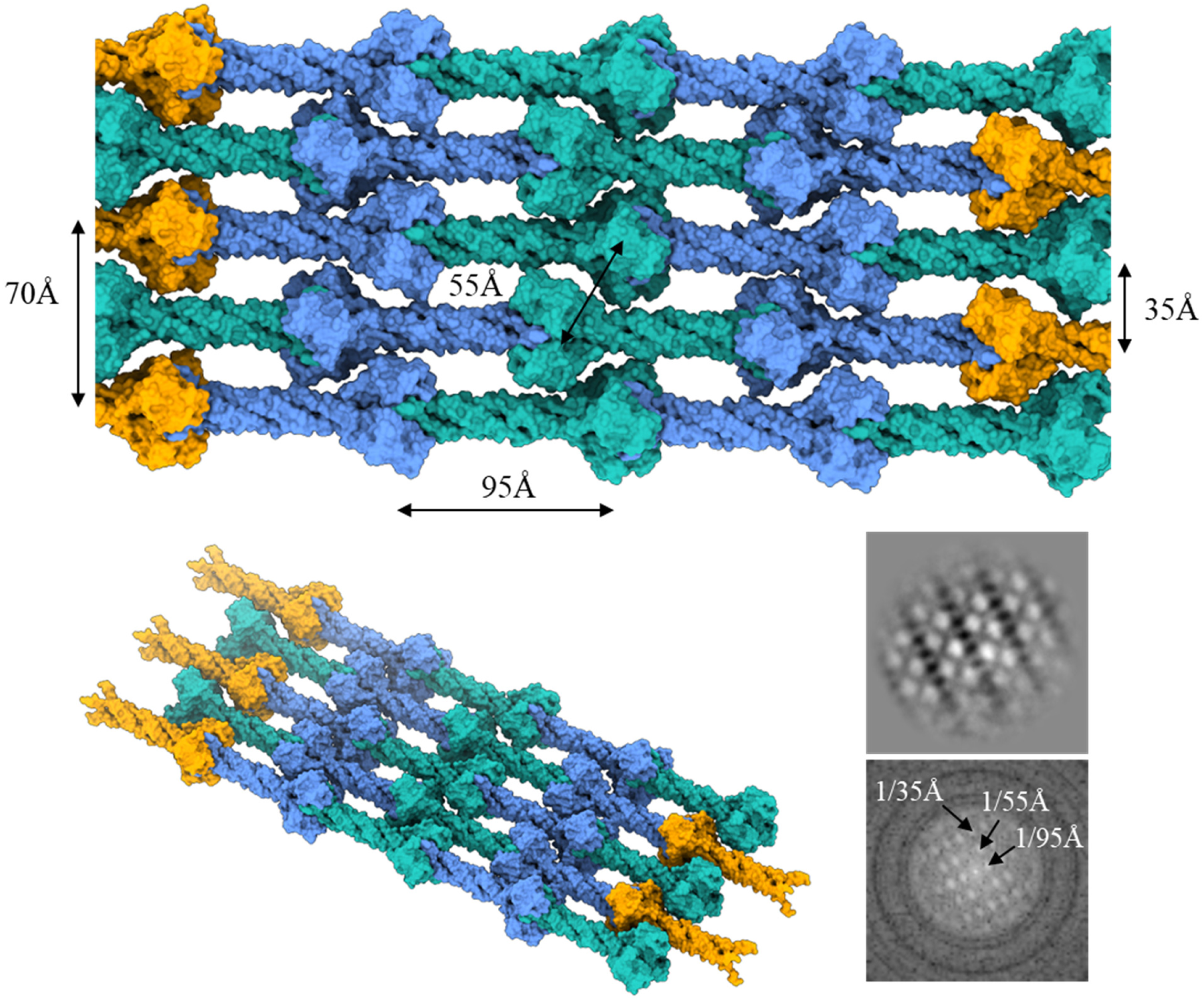
Molecular model of the hierarchical ultrastructure of F-ENA-2a. (a) and (b) Top and tilt views of an antiparallel crystalline array of F-ENA-2a fibers. (c) nsTEM 2D class average of F-ENA-2a fibers on a formvar supported copper mesh grid. (d) Power spectrum of the 2D class average in (c) with typical distances indicated.

**Supporting Figure 16.**
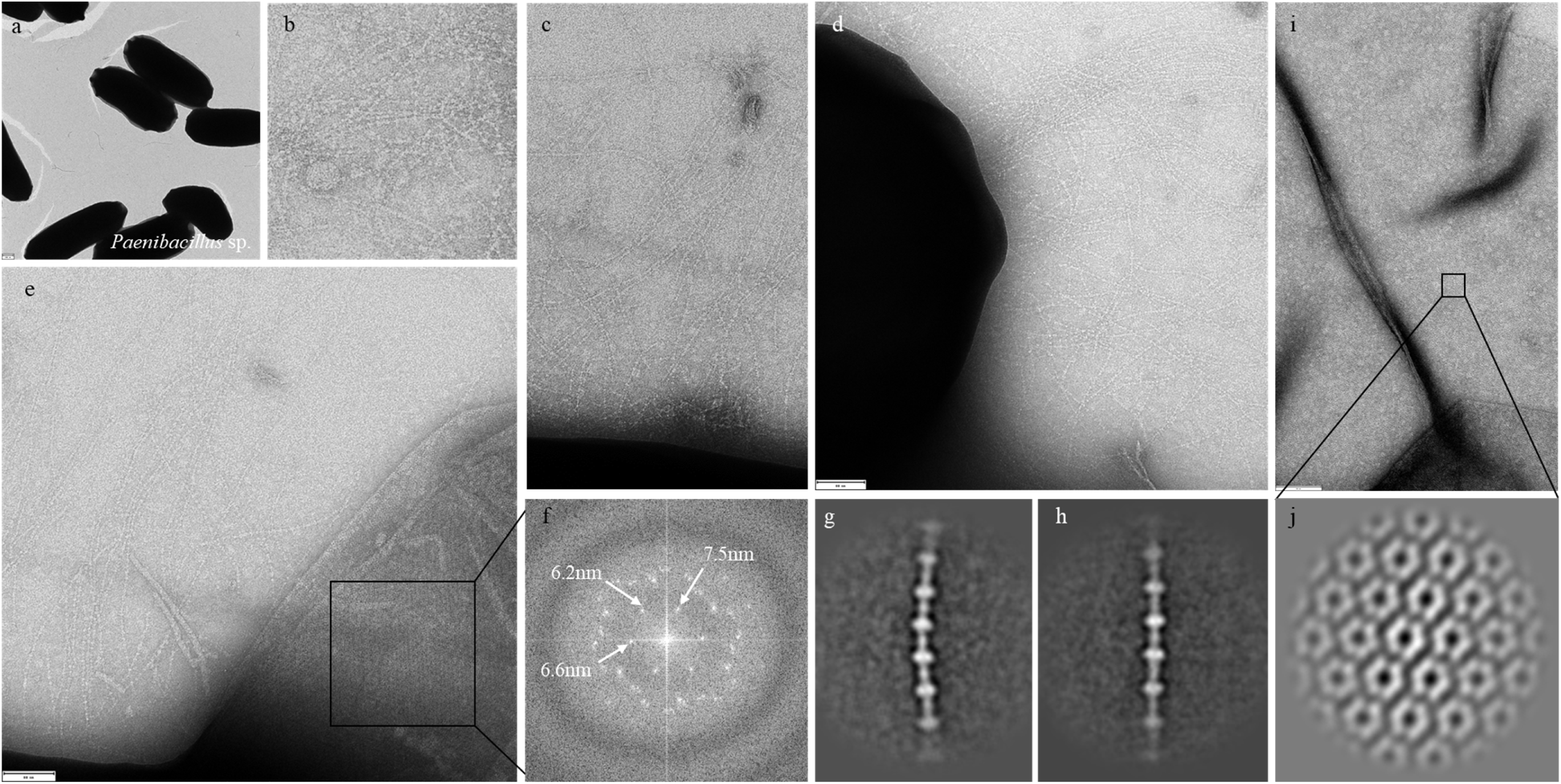
F-ENA-2a endospore appendages found on the surface of the spores of a *Paenibacillus* sp. strain in our collection. (a) Low magnification view of the spores. (b) – (c) High magnification views of the 3 nm diameter F-ENA-2a fibers, with typical ‘pearls-on-string’ morphology. (d) F-ENA-2a fibers seen emanating from the spore body. (e) nsTEM micrograph of exosporial segments with corresponding power spectrum shown in (f). (g) and (h) 2D class averages of F-ENA-2a fibers. (i) High magnification image of a single layered exosporium fragment with corresponding 2D class average (j) revealing a hexagonal array of hexameric units.

**Supporting Figure 17.**
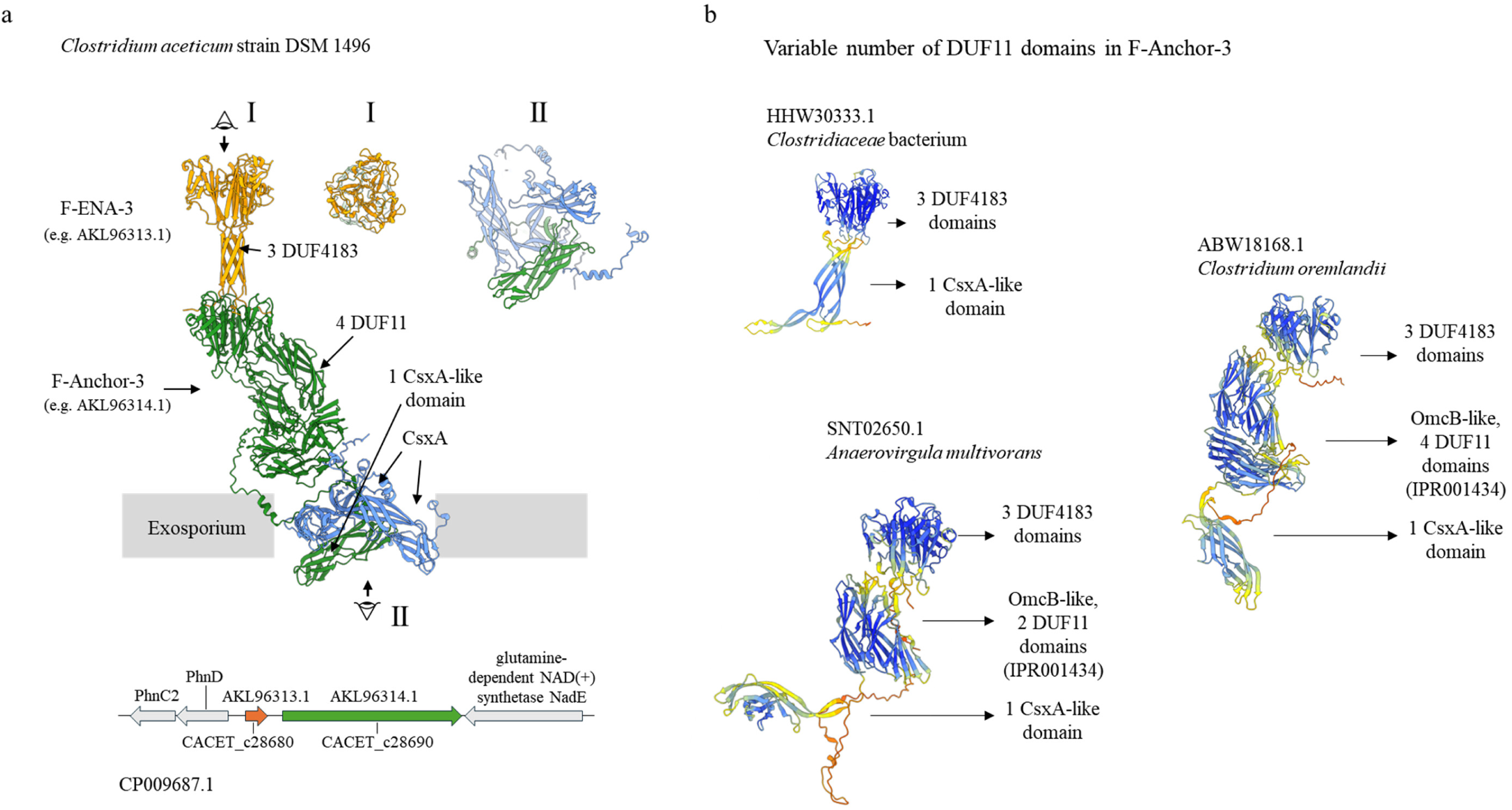
F-ENA anchoring mechanism in Clostridia. (a) AlphaFold3 prediction of a hetero-hexameric complex between 3 copies of F-ENA-3, one copy of F-Anchor-3 and two copies of CsxA; (b) genetic locus of F-ENA-3; (c) Genetic variation in F-Anchor-3 translates to variable number of DUF11 domains.

**Supporting Figure 18.**
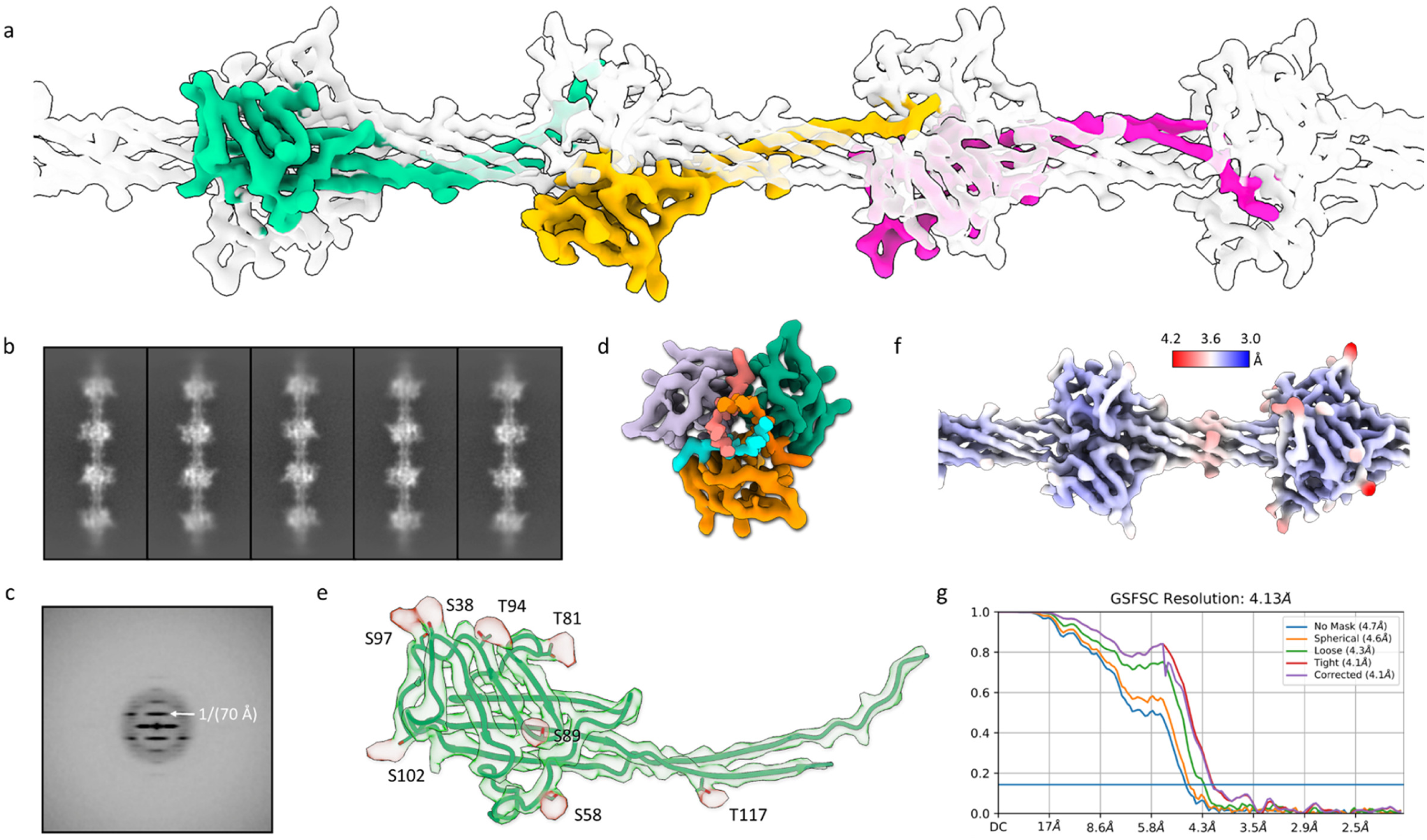
Helical reconstruction of F-type-3 ENA fiber. (a) Reconstructed cryo-EM volume of F-type-3 native endospore appendages (EMready sharpened); (b) 2D class averages: box size measures 346 Å vertically; (c) Average power spectrum of aligned particles; (d) Top view cross section; (e) atomic model docked in EMReady sharpened map with post-translational modification sites highlighted; (f) Surface colouring according to local resolution. The local-res map was generated by Local Filtering job in CryoSPARC; (g) The global resolution estimated by the map:map Fourier Shell Correlation.

## List of sequences

>F-ENA

MPIIQPFMASRRFTSTLGAGTGTGAAFAIAATACLNDAGTTATAFPTFTYYNLYVNGILQPSVNSSVTTGPTGAITIPGGDALDGGIPITIEFIVT*

>F-Anchor

MKRNDKLSINKGKIEPENIGPTFSALPPIYIPTGATGATGATGATGATGVTGPTGSTGPTGSTGSTGPTGSTGATGPTGATGETGATGATGATGETGATGATGATGETGATGVTGATGPTGATGITGITGETGETGATGETGATGETGATGPTGVTGETGATGATGPTGVTGATGATGVTGATGPTGGIGPITTTNLLFYTFSDGEKLIYTDADGIAQYGTTNILSPTEVSYINLFINGILQPQPFYEVSTGKLTLLDNQPPSQGASIILQFIIIN*

> F-Anchor_14-28_ (peptide)

NH_2_-IEPENIGPTFSALPPIYIPTG-COOH

>ExsF (crystallized construct)

MHHHHHHTLPAFGFAFNASAPQFASLFTPLLLPSVSPNPNIPVPVINDTVSVGDGIRILRAGIYQISYTLTISLDNSPVAPEAGRFFLSLGTPANIIPGSGTAVRSNVIGTGEVDVSSGVILINLNPGDLIQIVPVQLIGTVDIRAAALTVAQIS*

>F-BclA

MFINFSLLDINRIILPHSPTGATGSTGATGATGPTGATGSTGPTGPTGSTGPTGSTGSTGTTGATGATGATGATGATGSTGTTGSTGATGATGATGATGSTGTTGATGATGATGSTGTTGATGPTGATGSIGLTGSTGSTGTTGATGSTGPTGPTGSTGSTGSTGATGSTGPTGSTGATGSTGPTGSTGSTGTTGATGSTGPTGTTGATGSTGPTGPTGSTGTTGTTGATGATGATGSTGSTGPTGATGSTGLTGSTGTTGTTGTTGATGSTGPTGTTGSTGPTGATGSTGPAGPTGATGSTGSTGPTGTTGSTGATGATGSTGPAGPTGATGSTGATGPTGSTGTTGATGATGSTGLTGSTGTTGATGATGATGSTGLTGSTGSTGTTGATGATGSTGPTGATGSTGPTGATGATGSTGLTGSTGTTGATGATGSTGATGATGSTGPTGPTGSTGPTGATGPTGATGATGPTGPTGATGSTGPTGPTGATGSTGSTGPTGSTGSTGLTGSTGTTGTTGTTGATGSTGPTGTTGSTGATGATGATGPTGPTGSTGLTGSTGATGTTGTTGATGSTGPTGTTGSTGATGATGATGPTGPTGSTGATGATGATGPTGPTGATGATGATGSTGPTGTTGATGSTGLTGSTGATGSTGTTGATGATGATGATGATGATGATGATGATGATGATGATGATGATGATGATGSTGPTGFTGSTGPTGAASVGLTNYLYVFDTTNQSIAVGSSVTFNTNGPITGTALSHITGTGNIIINTLGTYVAEFQLQASRENQFSLELNGTPIPGGRFGTGSPHSINQGTAAFTVTVVPSTLTLINNTSSAGTITLSNSDGGSLTNVSASISIFQVG*

>F-BclA-CTD (crystallized construct)

MHHHHHHSSGENLYFQGSSGSVGLTNYLYVFDTTNQSIAVGSSVTFNTNGPITGTALSHITGTGNIIINTLGTYVAEFQLQASRENQFSLELNGTPIPGGRFGTGSPHSINQGTAAFTVTVVPSTLTLINNTSSAGTITLSNSDGGSLTNVSASISIFQVG*

>Ena2A (S-ENA)

MSCECSGSALTCCPDKNYVQDKVCSPWSGTVVATAITNVLYNNNINQNMIGTGFVRYDVGP APITLTVLDAAGATIDTQTLNPGTSIAFTYRRFVTIEVTLPAATAGTYQGEFCITTRYPLS*

>F-ENA-2a

MPVIKPVIVAVSSAPVSTGGVIATTVSPTVARFYAAITAAMIAGGVTTIPAASFLDDADAPVAALPVPAANGYYNVYINGILQQGGLSTLTAVSLALASGDFVEGTPVLLEVGTFGGDSTLTTQPTISAPTITIIS*

>F-Anchor-2

MPGQLGGRGPQGDLGPQGVQGVRGEVGPAGPTGPAGPIGQPGPAGPVGPAGTAGSEGPQGAAGATGATGATGATGPAGPMGETGAAGPAGPAGPPGPAGPLPAGIAVLPSSALYLAFMQEDTSGGPVTIDAGEFTVNGAPAAGFEGLGPNAYSMLYLNGVPQEQDLYALTSTAVTIDLDGSTLLAGTPVMVQIVSFSVEITA*

>F-ENA-3

MATTLFKLVMDAVTETETYTNPEVEKYFYNFDEADLDVDTLTIPATSFFDDAGDPVTSNLVTLADDNGYYLLFVNGALQQSSLFTVSTDGSEVVITQASTIPVGAPITLVVNNFAPTSTSTTTVTT*

>F-Anchor-3

MCKATPLKSKVYQYNALSDGKKRMYTNQDELLQYGNKGILDPHKVSYYSLFINGVLQPKTNYIIEKGLLFLKTKDLPLKGSPIIITFVSFIDKEILKLNTAIAEGSVPSGSIFHGPVTDVDIILEETVQSTALYLKLEKVITSGPAFIPTGHIAAWEFTLIVTNTVNMPISNIVVTARTLLDTTLNTTNLSLSQGNVGINHSIITWNVGILDVGESAIATFKLEGFFKADGVRFIDRAFAVGDTSLGTIKSSIASGKAIQVVKGLSITEAITSGSLNVVMKKNNKWRVEIKIVNLSDASISNILATDTLLIESIKSIEIVSLSQGSATIADNKILWKIDVLEGFRTAVLVVDIIGAFTIEGYRNLDSVMVVGIIASGEIFTGPSKDIRIVVSPNEKLPEDQLLLQKFITTEPLVAFLGKPRKWCFALKVINLTKDVLENLLVTDYILLEEFNDIHTLFVSSGDILIAHNTILWNVEELSPGETLTAVFEVNGFFNARGIRSLSRGVASGFNGNSATCTMSHMVSSPPIKVLDFIHNLKSSYILADKVYGQCRQQNCFEDITISIDNSNFKNILFKQGFIVKDTLVVTNMIDRPGFKRLQFLIRIPFEIITESGNRIKGYLPDLSEDIILFIPEARNEFPFNILVETSTKLLKEAARLNNQLTFTAGASIVMKVVGRVQLLLPTFEFSPQPPCCQEFNKNLICDIFQSRNFPDFFPGQSALNFHRKAARTNKTKQCPLIFGNLTIEKYITAGPLEVTTSAFNTWRMEIRISNDGYGPVSQVMMIDTLFLDHLTQINVLSLSQGNVSQEKDKILWDIGTLNSAATVVLIAEFTGFFNSKNKKPISTKTYQYNTISDGIKRVFTNDDELKMYGDEGILDPNEVSYFNLFINSVLQPQTNYIVKKGLLLLITIDVPPKGVPITLESIIIEDIHHQLLEVATYQYTAFGSGKKIYTNIDELRMYGNQGILDPRQISYQNLFINGVLQPPMNYVVEKGLLLLKTEDIPLEKSPISLQFITLFL*

>F-ENA-4

MATSLFKLVMSATTSTTTDTNPEVERYFYRLNPAQRTSGTLTIPSTSFTNSSGAAVTSNIVTAATDNGYYLLFINGAFQETSLFTVAASQVRITQTSLVPTSAPITLVVNNFAPTSTSTTTVRS

---

**Supplementary Table 1.**
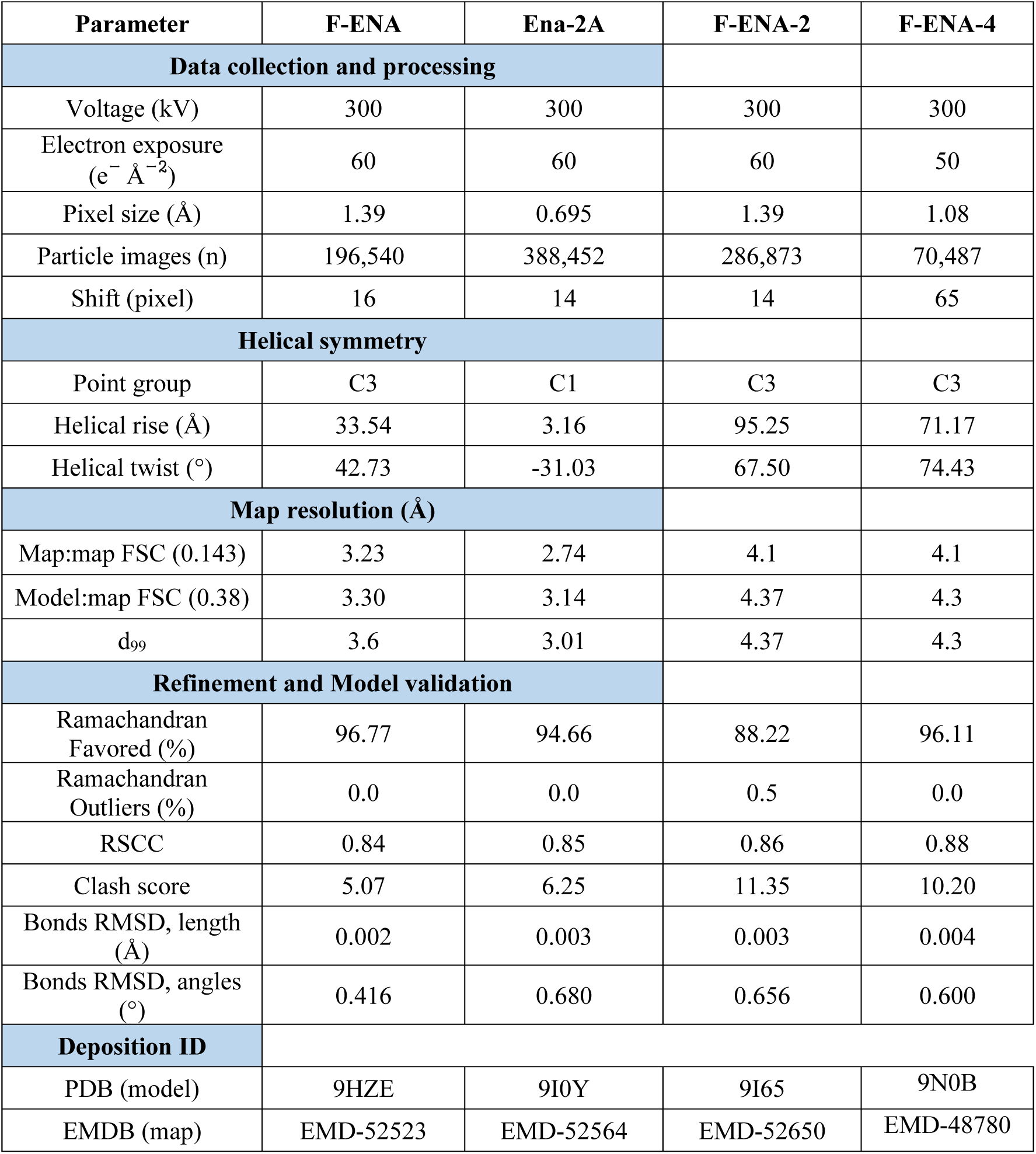
CryoEM data.

**Supplementary Table 2.** Crystallographic data.

|  | **F-BclA-CTD** | **ExsF/F-Anchor<sub>14-35</sub>** |
| --- | --- | --- |
| **Data collection** |  |  |
| Space group | P1 | P 21 |
| Cell dimensions |  |  |
| a, b, c (Å) | 71.8, 113.4, 113.5 | 51.29, 91.80, 99.93, |
| $\alpha$, $\beta$, $\gamma$ (°) | 60.11 89.19 86.35 | 90, 101.57, 90 |
| Resolution (Å)* | 49.235-2.292 | 50.25 - 1.922 |
| R<sub>pim</sub> | 0.096 (0.423) | 0.103 (0.969) |
| R<sub>merge</sub> | 0.148(0.617) | 0.232(1.965) |
| I/σ(I) | 4.9 (1.6) | 5.9 (0.8) |
| Completeness (%) | 97.8 (86.5) | 100 (97.6) |
| Multiplicity | 3.3 (2.9) | 6 (5) |
| **Refinement** |  |  |
| Resolution (Å) | 2.29 | 1.92 |
| Number of unique reflections | 128674 (8218) | 69027 (3362) |
| R-work/R-free | 0.185/0.242 | 0.184/0.215 |
| Number of atoms | 22780 | 7885 |
| protein | 20918 | 7144 |
| ligands/ions | 46 | 0 |
| water | 1816 | 741 |
| Average B-factors | 17.25 | 33.61 |
| protein | 17.20 | 33.10 |
| ligands/ions | 11.80 | / |
| water | 17.97 | 38.53 |
| RMS deviations |  |  |
| Bond angles (°) | 0.87 | 0.85 |
| Bond length (Å) | 0.007 | 0.007 |
| Ramachandran |  |  |
| favoured (%) | 97.02% | 96.66% |
| allowed (%) | 2.42% | 3.24% |
| outliers (%) | 0.56 % | 0.10 % |
| **PDB code** | **9H38** | **9H3D** |

\* Values in parenthesis refer to the highest recorded resolution shell.

## Materials and Methods

### Biofilm formation

Btk and *Paenibacillus* sp. were sporulated via nutrient starvation on solid LB agar medium. For this, an overnight LB culture of either strain was spread on a LB agar plate and grown to confluency over a period of 72h at 30°C.

### Negative stain transmission electron microscopy (nsTEM)

nsTEM imaging of bacterial biofilms was done using formvar/carbon-coated copper grids (Electron Microscopy Sciences) with a 400-hole mesh. The grids were glow-discharged (ELMO; Agar Scientific) with 4mA plasma current for 45 seconds. A section of a confluent microcolony was scraped from an LB agar plate and resuspended in Milli-Q. 3 μl of the bacterial spore suspension was applied onto the glow-discharged grids and left to adsorb for 1 minute. The solution was dry blotted, followed by three washes with 15 μl Milli-Q. Next, 15 μl drops of 2% (w/v) uranyl acetate were applied three times for 10 seconds, 2 seconds, and 1 minute respectively, with a blotting step in between each application. The excess uranyl acetate was then dry blotted with Whatman type 1 paper. All grids were screened with a 120 kV JEOL 1400 microscope equipped with LaB6 filament and TVIPS F416 CCD camera.

### *Ex vivo* Isolation of S-type and F-type endospore appendages from Btk

To dislodge the ENA fibers from the *Bacillus thuringiensis* subsp kurstaki spore surface, 500 µl of the bacterial spore suspension was subjected to sonication on ice using a Qsonica Q500 sonicator equipped with a 1/16” microtip probe using the following settings: 15/30 sec on/off, 2 min total time, 30% amplitude. Following sonication, the spore suspension was loaded onto a 1.8mL 60% (w/v) sucrose cushion and centrifuged at 20.000 rcf for 1hour. The cleared supernatant was carefully pipetted off the sucrose layer and subjected to 3 rounds of washing in Mili-Q via repeated centrifugation (1 h, 20.000 rcf) and resuspension of the pellet. nsTEM imaging of the resuspended pellet confirmed the presence and enrichment of the S- and F-type ENA fibers.

### Cloning of ENA2A, F-ENA-2a, F-BclA-CTD and ExsF

All genes were in cloned in the pASK-IBA3^+^ vector via Gibson assembly (New England Biolabs). The genes for recombinant expression of ENA2A (WP_001277547.1), F-ENA-2a (A0A1I1C8X4) and the C-terminal domain of F-BclA (F-BclA-CTD; BTK_RS34675 residues 721 to 863) were ordered as double-stranded synthetic DNA constructs (gblocks Gene Fragment, IDT) with the appropriate overhangs, while the *exsF* gene, in which the first 23 residues were replaced by a 6xHis tag, was amplified from the *B. paranthracis* genome. Gibson assembly mixtures were transformed into *E. coli* DH5a cells and plated on LB agar plates supplemented with 100 µg/ml ampicillin. Positive clones were identified via colony-PCR followed by Sanger-sequencing (Mix2seq, Eurofins).

### Recombinant production of S-ENA and F-ENA-2a fibers

Recombinant S-ENA and F-ENA-2 fibers were produced by heterologous expression in the cytoplasm of *Escherichia coli*. A single colony of DH5a(pASK-IBA3^+^-*ena2A*) or DH5a(pASK-IBA3^+^-*f-ena2A*) was grown overnight in LB supplemented with 100 µg/ml ampicillin. The overnight preculture was diluted 50x fold in fresh LB supplemented with 100 µg/ml ampicillin and grown to exponential phase at 37°C, shaking at 180 rpm. Recombinant expression was induced by adding 200 µg/l final anhydrotetracycline at an OD_600nm_ of 0.8 and incubated at 20°C, 180 rpm for 3 hours. Cells were pelleted at 4000 rcf and stored at −20°C until further use.

To isolate S-ENA fibers from the cytoplasm of *E.coli*, stored pellets were resuspended in 50 mM Tris-HCl pH 7.5, 250 mM NaCl, 10 mM Ethylenediaminetetraacetic Acid (EDTA), 0.5 mg/ml Henn egg white lysozyme (HEWL) (Merck) and continuously stirred at 37°C for 1 h. Next, sodium dodecyl sulfate (SDS) was added to the lysate to a final concentration of 2% (w/v), and the suspension was heated to 95°C for 1h. The cleared lysate was centrifuged for 1 h at 35.000 rcf, and the pellet was resuspended in Mili-Q. This process was repeated threefold to remove residual SDS and obtain a pure suspension of S-ENA fibers. To isolate F-ENA-2a fibers from the cytoplasm of *E.coli*, stored pellets were resuspended in 50 mM Tris-HCl pH 7.5, 250 mM NaCl, 10 mM EDTA, 0.5 mg/ml HEWL, 1% (w/v) n-Dodecyl-b-D-maltopyranoside (DDM) and continuously stirred at 37°C degrees for 1 h. The lysate was centrifuged for 1 h at 35.000 rcf, and the pellet was resuspended in Mili-Q. This process was repeated threefold to remove residual DDM and obtain a pure suspension of F-ENA fibers.

### Recombinant production and purification of ExsF and F-BclA-CTD

Recombinant ExsF and F-BclA-CTD were produced by heterologous expression in the cytoplasm of *Escherichia coli*. A single colony of DH5a(pASK-IBA3^+^-*exsF*) or DH5a(pASK-IBA3^+^-*f-bclA-ctd*) was grown overnight in LB supplemented with 100 µg/ml ampicillin. The overnight preculture was diluted 50x fold in fresh LB supplemented with 100 µg/ml ampicillin and grown to exponential phase at 37°C, shaking at 180 rpm. Recombinant expression was induced by adding 200 µg/l final anhydrotetracycline at an OD_600nm_ of 0.8 and incubated at 20°C, 180 rpm for 3 hours. Cells were pelleted at 4000 rcf and stored at −20°C until further use.

Stored cell pellets were thawed and resuspended in 100 mM Tris pH 7.5, 500 mM NaCl, 10 mM EDTA, 0.5 mg/ml HEWL, cOmplete Protease Inhibitor Cocktail (Merck), incubated for 30 min at 37°C and passed three times through a Microfluidics LM10 Microfluidizer operated at 20 kpsi. The lysate was centrifuged for 45 min at 35.000 rcf to pellet remaining cell debris. The cleared supernatant was supplemented with 20 mM imidazole pH 8.0 final concentration and loaded onto a 5mL Histrap High Performance prepacked column (Cytiva) that was equilibrated with 5 column volumes (CV) of 100 mM Tris pH 7.5, 500 mM NaCl and 20 mM imidazole (Buffer A). Following sample loading, the column was washed with 10 CV of Buffer A, and gradient (flow rate 2.5 ml/min; 50% A/B gradient over 30min) eluted with 100mM Tris pH 7.5, 150mM NaCl and 500mM imidazole (Buffer B) whilst fractionating the eluate. SDS-PAGE analysis was used to monitor sample purity. Pure F-BclA-CTD fractions were pooled and dialyzed against 25 mM Tris pH 7.45, 500 mM NaCl using Spectra/por 3 Dialysis Membrane, MWCO 3500 Spectrum and concentrated using a 10 kDa MWCO Amicon Ultra Centrifugal Filter, to a final concentration of 18.4 mg/ml. The protein concentration was determined based on absorption at 280 nm using a NanoDrop One UV-Vis spectrophotometer, an extinction coefficient of 5960 M^-1^ cm^-1^ and a molecular weight of 16583.22 Dalton.

### Crystallization, X-ray data collection, processing and structure determination of F-BclA-CTD and ExsF/F-Anchor_14-35_

Protein crystallization screens were setup using the sitting drop vapor diffusion method and a Mosquito nanoliter-dispensing crystallization robot (SPTlabtech). For F-BclA-CTD, plate-like crystals appeared within one week in a condition C2 of the JCSG-plus HT-96 crystallization screen (Molecular Dimensions) containing 1M LiCl, 0.1M citrate pH 4.0, 20% (w/v) PEG 6000. ExsF/F-Anchor**_14-35_** crystallized in 5 % v/v T-mate pH 7, 0.1 M sodium cacodylate pH5.3, 15 % v/v PEG smear broad and 10 %v/v ethylene glycol (BCS screen). Crystallization was further optimized by manual setup of replicated crystallization conditions in hanging drop geometry using a Greiner Bio-One ComboPlate 24-well Protein Crystallization Plate, Pre-greased and crystals appeared within one week. The crystallization buffer was supplemented with 10% glycerol, and crystals were mounted in nylon loops and flash-cooled in liquid nitrogen. X-ray diffraction data were collected at 100 K using the Beamline Proxima 2 (wavelength = 0. 9801 Å) at the Soleil synchrotron (Gif-sur-Yvette, France). For F-BclA-CTD, diffractograms were processed using Autoproc^36^ and Staraniso^37^ at 2.29Å in P321 with unit-cell dimensions *a* = *b* = 112.88 Å, *c* = 111.71 Å, *a* = *b* = 90.0° and *g* = 120.0°. ExsF/F-Anchor_14-35_ diffracted to 1.92 Å and was processed using AutoProc in P 21. Both crystal structures were determined by molecular replacement using phaser from the phenix suite^38^ and using the AlphaFold2^39^ predictions of the F-BclA-CTD or ExsF as search models. The structures were refined through iterative cycles of manual model building with COOT^40^ and reciprocal space refinement with phenix.refine to R values of Rwork/Rfree of 0.18/0.24 for F-BclA-CTD and 0.18/0.21 for ExsF/F-Anchor_14-35_, respectively. The crystallographic statistics are shown in Supplementary Table 1. Structural Figures were generated with ChimeraX^41,42^. Atomic coordinates and structure factors have been deposited in the protein data bank^43^ (PDB) under the accession codes 9H38 (F-BclA-CTD) and 9H3D (ExsF/F-Anchor_14-35_).

### Cryo-electron transmission microscopy

Quantifoil® holey Cu 400 mesh grids with 2-µm holes and 1-µm spacing were glow discharged in vacuum using plasma current of 5 mA for 1 minute (ELMO; Agar Scientific). For cryo-plunging, 3 µl of a fiber suspension was applied on freshly glow-discharged grids at 100% humidity and room temperature in a Gatan CP3 cryo-plunger. After 10s of absorption, the grid was machine-blotted with Whatman grade 2 filter paper for 3.5 seconds from both sides and plunge frozen into liquid ethane at −176°C. Grids were stored in liquid nitrogen until further use. In total, three datasets were collected: one for the *ex vivo* ENA fibers purified from Btk (dataset 1: 27,272 movies), one for recombinant Ena2A (dataset 2: 4,348) and one for recombinant F-ENA-2 fibers (dataset 3: 8,715 movies). High-resolution cryoEM movies were recorded using SerialEM 3.0.850 on a JEOL CRYO ARM 300 microscope equipped with an omega energy filter (operated at slit width of 12 eV). The movies were captured with a K3 direct electron detector run in counting mode at a magnification of 60K with a calibrated pixel size of 0.70 Å/pixel, and a total exposure of 60 e/Å2 over 60 frames.

Movies were imported into cryoSPARC v4.6.2^44^ for further processing. Movies were motion-corrected using Patch Motion Correction and defocus values were determined using Patch CTF. Exposures were curated and segments were picked using the filament tracer and extracted at 4x binning. After several rounds of 2D classification, initial estimates of the helical rise and twist values were obtained based on the 2D class averages (F-ENA; F-ENA-2) or were taken from the S-ENA structure (7A02), and were used in subsequent helical refinement jobs. Next, particles were reextracted at native unbinned resolution (Ena2A: 0.695 Å/pixel) or twofold binned (F-ENA/F-ENA-2: 1.39 Å/pixel) and used as input for a second round of helical refinement, followed by local and global CTF refinement, and helical refinement. In the end 196,540, 388,452, and 286,873 fibril segments were used for F-ENA, Ena2A and F-ENA-2, respectively. The resulting high-resolution volumes and particle stacks were used for reference-based motion correction after particle duplicate removal followed by a final helical refinement. Initial atomic models were built using either ModelAngelo^17^ without providing an input sequence for F-ENA (build_no_seq), and with providing input sequences for recombinant Ena2A. For F-ENA-2, an amber-relaxed AF2 model was generated of an F-ENA-2 hexamer using ColabFold^45^. Modelangelo generated models of F-ENA and Ena2A were real-space refined in Coot.9.8.95^40^ and phenix.refine (default settings with NCS constraints (90,91). The AF2 F-ENA-2 model was fitted into the reconstructed helical volume, and real-space refined in phenix.refine with tight restraints (target bonds rmsd: 0.05; Target angles rmsd: 0.5). Figures were created using ChimeraX 1.8.92^41,42^. Map and model statistics are found in Supplementary Table 1.

### Cryo-EM and modeling of F-type-3 ENA fibers from class Clostridia

Initially, spore-like architecture and endospore appendages were observed in an anaerobic, Gram-negative culture of *Syntrophus aciditrophicus* (DSM 26646), purchased from the DSMZ. Since recent advancements in cryo-EM and deep learning modelling tools now routinely allow for direct protein identification from cryo-EM maps at resolutions of 4 Å or better^17^, we proceeded directly to high-resolution cryo-EM imaging. This putative endospore culture (∼ 4.5 μl) was applied to glow-discharged, lacey carbon grids and plunge-frozen using an EM GP Plunge Freezer (Leica). Cryo-EM data were collected on a 300 keV Titan Krios equipped with a K3 camera (University of Virginia) at a pixel size of 1.08 Å/pixel and a total dose of approximately 50 e/Å². Patch motion correction and CTF estimation were performed using cryoSPARC^44^. Particles were then automatically selected using the “Filament Tracer” tool. These particles were subjected to multiple rounds of 2D classification to remove low-quality particles. Following this, 70,487 particles remained in the F-ENA-4 dataset, with a shift of 65 pixels between adjacent boxes. Possible helical symmetries were calculated from an averaged power spectrum generated from aligned, raw particles. Subsequently, 3D reconstruction was performed using “Helical Refinement,” testing all potential helical symmetries. The resolution of the final reconstruction was estimated using map:map FSC, model:map FSC, and d_99_. The final volume was sharpened with EMready^24^, and the corresponding statistics are listed in Supplementary Table 1.

The final helical reconstruction of the F-ENA-4 fiber achieved a resolution of 4.1 Å, as estimated by map:map FSC. To identify the protein sequence, ModelAngelo^17^ was used to build the protein backbone and predict amino acid sequences based on side-chain densities. A BLASTP search using this sequence against all proteins in GenBank yielded a top hit of MDF2656102, a protein encoded by an uncultivated member of the family Anaerovoracaceae within the class Clostridia, with 62% sequence identity. The second-best match had only 48% identity. To validate the appendage sequence, we designed several primer pairs targeting regions near F-ENA-4 and successfully amplified the gene from the spore genome, confirming that MDF2656102 is indeed the correct sequence. For the final model building, an AlphaFold prediction of MDF2656102 was docked into the cryo-EM map. The single subunit was manually refined in Coot^40^, the helical symmetry was applied to the model, and the resulting helical model was real-space refined in PHENIX^38^. Refinement statistics for F-ENA-4 are presented in Supplementary Table 1.

### Generation of deletion strains

Appendage-depleted mutants were constructed as described in Pradhan et al.^14^ using markerless gene replacement^46^ using Bt407 as a background strain. Gene replacement constructs, containing the start and stop codons of the target genes flanked by upstream and downstream homologous sequences were amplified with chromosomal DNA as the PCR template. The homologous sequence were ligated into pMAD vectors and cloned into One Shot TOP10 chemically competent *E. coli* (Invitrogen, Thermo Fisher Scientific, USA) for plasmid amplification. Verification of the construct was performed through sequencing (LightRun, Eurofins, Luxembourg). The construct was passed through One Shot INV110 *E. coli* (Thermo Fisher Scientific) to demethylate the vector before transformation into electrocompetent Bt407 cells. Colonies containing the chromosomally integrated pMAD construct were identified by PCR and subsequently transformed with a demethylated pBKJ223 plasmid. The pBKJ223 plasmid encodes an enzyme that introduces a double-stranded DNA break in the integrated pMAD plasmid, leading to its excision from the chromosome and leaving behind a markerless gene deletion. Verification of the deletion was conducted using PCR with primers targeting regions upstream and downstream of the gene of interest.

### Determination of the spore versus cell ratio

A confluent macrocolony of Btk grown on agar was resuspended in miliQ water to a final OD_600_ of 34. 5µl of the suspension was deposited onto a microscope slide, sealed with a coverslip and imaged using an inverted microscope Leica DMi8 equipped with a DFC7000 GT camera using a 100x oil objective in phase-contrast mode. A total of 50+ image tiles were collected using LasX and stitched together into a single tiff file covering 374 x 238 cm^2^. The total number of spores and cells was determined using Fiji^47^, yielding 2878 spores and 179 cells, leading to a spore fraction of 94%.

### Sedimentation assays

For sedimentation analysis, spores of Bt407 and isogenic appendage-depleted mutants were prepared by streaking freshly cultured cells onto LB agar plates and incubating at 30 °C for two weeks or until sporulation reached ∼98%, determined by phase-contrast (PH) microscopy. Mature spores were scraped off the agar, washed three times in distilled water, and resuspended in sterile distilled water. Purity was verified via PH microscopy.

Spore suspensions (OD_600_ = 10) were mixed by vortexing for 15 seconds, placed in borosilicate tubes (14mm × 130mm, DWK Life Sciences), and left to settle. For each strain, a total of three spore batches were tested. A SONY ILCE-5100 camera with a SELP1650 lens captured images at 0, 3, 5, and 10 hours, then every 24 hours until full sedimentation. Images were analyzed in ImageJ 1.54f. Pixel intensity profiles of sedimenting spores were normalized to the highest intensity and converted to 200 datapoints using a randomized algorithm in RStudio (Version 2023.12.1). The profiles were plotted over time, with datapoints crossing 50% intensity extracted and plotted as a function of time.

### Phylogenetic Analysis

To probe for the presence of F-ENA across the Firmicutes phylum, all publicly available Genbank genomes (9342 genomes) belonging to the Bacillus/Clostridium group (taxid 1239; assembly-level complete) were downloaded from the NCBI database. Homo- and orthologs of F-ENA were searched in all assemblies with hmmsearch^48^ (881 genomes) using a hidden Markov model that was generated with hmmbuild using a manually curated multiple sequence alignment of F-ENA sequences obtained from a blastp^49^ search using WP_001121647.1 as a query. The inclusion threshold for hmmsearch was set to an E-value of 1e-7. The phylogenetic tree of the Firmicutes phylum was generated using phyloTv2 (https://phylot.biobyte.de/) using the NCBI taxonomy IDs of the corresponding assemblies as node identifiers, and imported into iTOL^50^ for visualization using the online server (https://itol.embl.de/).

For the more detailed search of F-ENA homo- and orthologs across the *Bacillus cereus* group sensu lato, the BtyperDB v1 database (5976 genomes)^51^ was downloaded from https://www.btyper.app/, and hmmsearch was run against all assemblies with an E-value threshold of 1e-7 (5647 genomes). The phylogenetic tree described in Ramnath *et al*^51^ and the corresponding meta-data table was kindly provided by Laura M. Caroll. Tree figures were generated using https://microreact.org/^52^.

## Author contributions

M.S. and H.R. designed the project. M.S. performed cryogenic freezing, nsTEM and cryo-EM imaging and data processing. U.L.J. and M.A. designed and generated the knockout strains, and performed sedimentation assays. A.S. and L.D.T. performed protein purification, crystallization and crystallographic analysis. M.F. supported the EM work. N.V.G., A.S. and I.V.M. performed cloning and protein expression. N.F.B., D.P.B., E.H.E., M.K. and F.W. performed the sample and image analysis for F-type-3 ENA fibers from the class Clostridia. M.S. and N.V.G. wrote the manuscript with contributions from all authors.

## Data Availability Statement

The *ex vivo* F-Ena cryo-EM map and the corresponding atomic model were deposited to the EMDB and the PDB under entries 9HZE and EMD-52523 respectively. The recombinant Ena2A cryo-EM map and the corresponding atomic model were deposited to the EMDB and the PDB under entries 9I0Y and EMD-52564, respectively. The recombinant F-ENA-2 cryo-EM map and the corresponding atomic model were deposited to the EMDB and the PDB under entries 9I65 and EMD-52650 respectively. The recombinant F-ENA-4 cryo-EM map and the corresponding atomic model were deposited to the EMDB and the PDB under entries 9N0B and EMD-48780, respectively. The crystal structures of ExsF/F-Anchor_14-28_ and F-BCLA-CTD were deposited to the PDB under entries 9H3D and 9H38, respectively.

## Acknowledgements

We thank Dirk Reiter at the VIB-VUB Facility for Bio Electron Cryogenic Microscopy (BECM). We thank Dr. Tom Delmont for his helpful discussion. This work was funded by VIB, NMBU, FWO through grant G043021N to M.S. A.S was supported by the EMBO (ALTF-709-2021) and the Marie Skłodowska-Curie Actions (MSCA; SLYDIV project). M.A. recognizes the Grant from the Norwegian research council (NFR): 335029 - FORSKER22. EHE was supported by NIH GM122510.

